# Neurons, Muscles, and Venom: Elucidating a Neural-to-Secretory Pathway in Cephalopod Predation

**DOI:** 10.1101/2025.06.15.659772

**Authors:** Kim N. Kirchhoff, José Ramón Pardos-Blas, Amy Courtney, Belkes Stambouli, Andrew L. Sugarman, Daniel J. Vanselow, Melvin Williams, Christian Carrasco, Mariam Gelashvilli, Cecilia Cuddy, Praveena Naidu, William R. Schafer, Keith C. Cheng, Eve Seuntjens, Mandë Holford

**Affiliations:** Department of Organismic and Evolutionary Biology, Harvard University, Cambridge, MA, USA; Department of Chemistry, Hunter College, CUNY, NY, USA; Departamento de Biodiversidad y Conservación, Real Jardin Botanico, CSIC, Madrid, Spain; MRC Laboratory of Molecular Biology, Cambridge, United Kingdom; Department of Pathology, Pennsylvania State University College of Medicine, Hershey, PA, USA; Laboratory of Molecular Biology, The Rockefeller University, NYC, NY, USA; Programs in Biology, Biochemistry, and Chemistry at CUNY Graduate Center, NYC, NY, USA; Institute for Computational and Data Sciences, Pennsylvania State University, University Park, PA, USA; Department of Biology, KU Leuven, Leuven, Belgium; Department of Invertebrate Zoology, The American Museum of Natural History, NYC, NY, USA

**Keywords:** Cephalopods, Venom gland, Neuronal control, Secretion

## Abstract

Venom plays a central role in the predatory ecology of coleoid cephalopods (octopuses, squids, and cuttlefish), yet the mechanisms governing venom release from the posterior salivary gland (PSG) are unknown. Using a multimodal approach combining X-ray micro-histotomography, histological stainings, *in situ* hybridization, comparative phylogenetics, and *ex vivo* imaging across multiple coleoid species, we characterize the structural and neuronal regulatory organization of the PSG. We verified that the gland comprises two distinct tubular systems: secretory tubules specialized for venom production and smooth–striated tubules positioned to facilitate venom transport toward the beak for injection into its prey. Molecular localization of filamentous and α-actin confirms a circular smooth muscle layer surrounding the tubules. Mapping of six neuronal markers, including neurofilament (NF-H), synapsin, and muscle-type nicotinic acetylcholine receptors, reveals dense and stereotyped neural innervation closely associated with the muscular compartments. Comparative phylogenetic analyses of cys-loop ligand-gated ion channel sequences indicate a predominancy of excitatory acetylcholine- and dopamine-gated receptors in coleoid venom glands, implicating potential molecular agents involved in neural control of venom release. Consistent with neural regulation, *ex vivo* stimulation of the PSG elicits calcium signaling throughout the gland. Together, our results reveal a conserved venom gland structure among octopuses, squids, and cuttlefish, with a spatially distinct neuromuscular tissue organization indicating venom production and release sites modulated by a network of neuronal agents. This work provides a mechanistic framework into venom gland organization and molecular regulation of venom release in one of the oldest venomous lineages.

## Introduction

Mollusks are the second-largest animal phylum, within which cephalopods (approximately 800 species) stand out for their complex cognition, camouflage, and highly developed central brain (1,2). Although these traits have attracted substantial research attention, cephalopod’s use of venom for predation, defense and sexual competition is understudied. Venoms are complex biochemical cocktails—often composed of small molecules, peptides, and proteins—that have convergently evolved across major animal lineages (3,4) and have demonstrated therapeutic relevance (5). Despite the prevalence of venom in nature, the mechanisms underlying its biological processes such as production, regulation and release are not well understood.

Marine venomous organisms like cnidarians (*e*.*g*., jellyfish) and coleoid cephalopods (octopuses, squids, cuttlefish) comprise some of the oldest venomous creatures dating back 500 million years (6) (Fig. 1a). Their evolutionary history, the wide presence of a venom system, the remarkable diversity of niche adaptations and the husbandry and breeding advantages of some species convert cephalopods into an ideal model for venom assessment in marine invertebrates. However, this research area is still pioneering and number of characterization studies is limited. The current state of the art is that a coleoid’s specialized venom system consists of the posterior salivary gland (PSG), which can be single (in myopsid squids) or paired (in sepiolid squids, cuttlefish and octopuses), and is hypothesized to be the primary venom producing tissue (Fig. 1b). In the PSG, venom is stored in secretory granules that are released to the salivary papilla via the salivary duct, ultimately entering the buccal mass to enable immobilization of the prey (7,8) (Fig. 1b-e and Fig. S1). However, the mechanism(s) controlling venom secretion remains unanswered. Prior morphological and histological studies in octopus from Young in the 1960’s suggest venom release is controlled by dual innervation of the PSG (7,9). Specifically, it has been proposed that the superior buccal lobe innervates the basal membrane in the secretory tubules of the gland, while the fibers of the subradular ganglion regulate the smooth muscles in the gland and those associated with the salivary duct and papilla (7,9,10). However, molecular aspects of the venom release mechanism characterizing neuronal subtypes, neurotransmitter systems and signaling pathways have not been identified in any single species or in a broader comparative framework across the phylogenetic tree of cephalopods.

**Figure 1.**
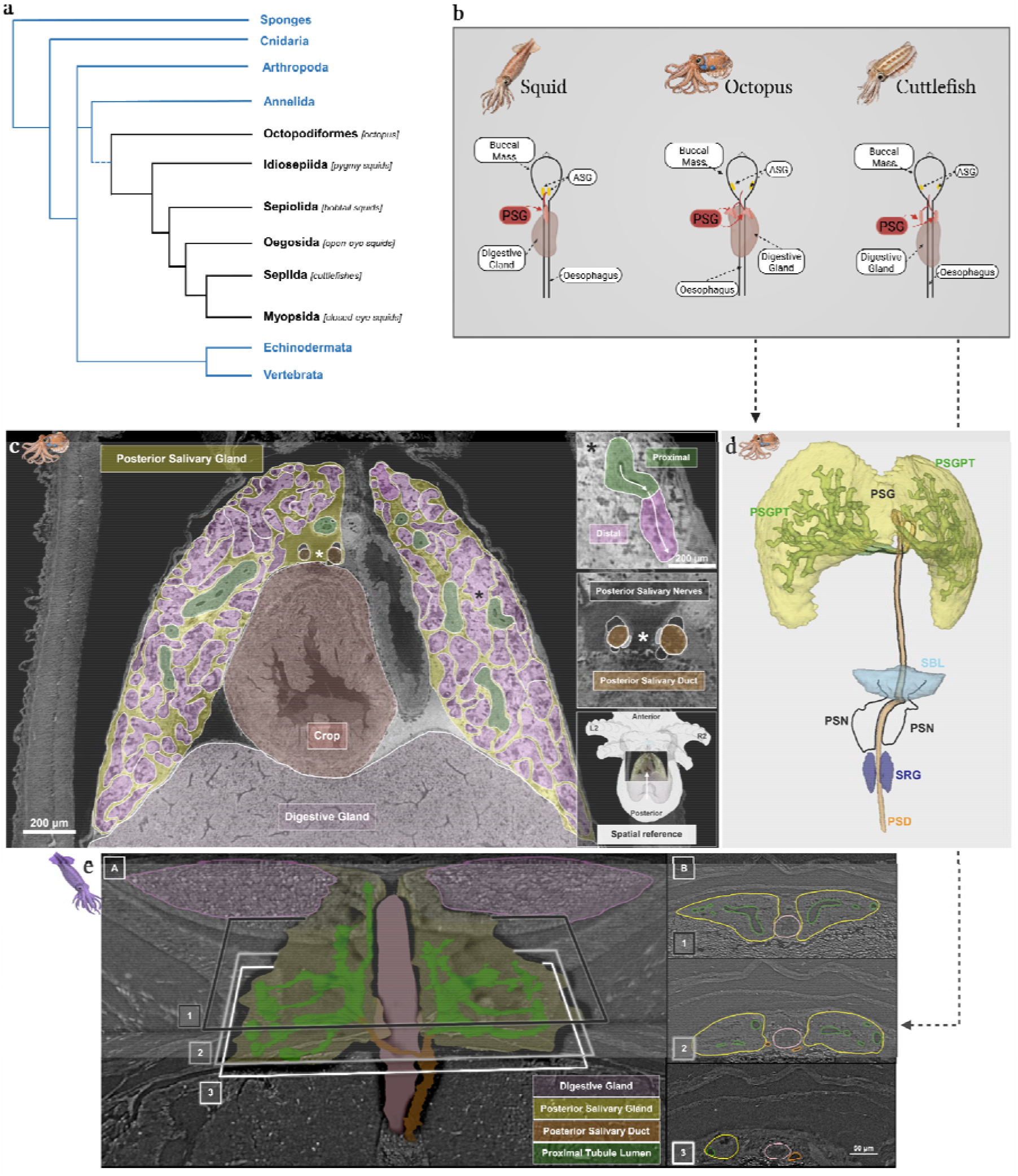
Cephalopods produce venom in their posterior salivary glands, which can be paired or single. **a**, Phylogenetic tree of venomous animal taxa highlighting the position of venomous coleoid cephalopods, *i*.*e*. octopuses, squids, and cuttlefish (adapted from (11,12). **b**, Anatomy of the venom system in coleoid lineages comprising the posterior salivary gland (PSG), *i*.*e*. venom gland (single or paired), venom duct, salivary papilla and buccal mass (dorsal view; adapted from (13) in BioRender https://BioRender.com). **c**, Micro-computed tomography (CT) 2D reconstruction of a transversal single 4 µm section of an *Octopus bimaculoides* hatchling shows the paired PSGs, proximal-striated (green) and distal-secretory (pink) tubules, posterior salivary nerves (black), posterior salivary duct (PSD) and digestive gland (purple) in corresponding color code to the 3D model in d. **d**, Micro-CT based 3D reconstruction highlights the segmentation of the gland’s tubular system (PSGT; green) as well as neuronal tracing from the posterior salivary nerve fibers (PSN; black) of the superior buccal lobe (SBL; light blue) to the PSG (yellow) in *O. bimaculoides* hatchling. The subradular ganglion (SRG; dark blue) and PSD are also shown. **e**, Micro-CT 2D reconstruction of a transversal single 4 µm section of an *Euprymna berryi* juvenile shows the paired PSGs, proximal-striated (green), PSD and digestive gland (purple).

Here, we address these gaps in knowledge of venom release through comprehensive histological profiling of the neuronal and muscular molecular organization of PSGs across all three venomous coleoid cephalopod lineages: octopus - the California two-spot octopus (*Octopus bimaculoides*), squid - the longfin inshore squid (*Doryteuthis pealeii*, myopsid squid) and the hummingbird bobtail squid (*Euprymna berryi*, sepiolid squid), and cuttlefish - the dwarf cuttlefish (*Ascarosepion bandense*) and the common cuttlefish (*Sepia officinalis*) (Fig. 1a,b, Table S1). We use tissue-specific markers and *ex vivo* live staining recordings to identify key molecular components of muscular and neuronal tissue, analyze their distribution patterns from pre-to postsynaptic sites, and integrate them into a mapping of the potential neuronal signaling pathway. Additionally, we performed a phylogenetic analysis for a key ligand-gated ion channel family involved in synaptic transmission across metazoan taxa to identify the presence of homologous channels in the PSG of cephalopods.

By integrating structural, cellular, neuromuscular signals and channel homology between the more studied octopus (Octopodidae) and squids and cuttlefish (loliginid/sepiolid), we provide the first cross-species integrative comparison and neuromuscular characterization of coleoid venom glands. Our findings allow insights into the nerve subtypes and neurotransmitters that may regulate coleoid venom release, elucidating spatial information about their neuromuscular structures and communication sides that have not been previously experimentally identified.

## Results

### Conserved *venom gland structure across coleoid lineages*

To determine if there are differences in the venom secreting tissue across coleoids, we performed morphological characterization of the venom gland (PSGs) in octopus (*O. bimaculoides*), squid (*E. berryi*) and cuttlefish (*A. bandense*) specimens. Tissue micro-computed tomography (micro-CT, histotomography) has been shown to be a powerful tool to reconstruct tissue structures, sensory pathways and neuronal connections in octopuses (14). The micro-CT reconstructions of the PSG in an *O. bimaculoides* hatchling indicate two differentiated tubule types distinguished by morphology and clear differences in texture (Fig. S2). The two types present correspond to secretory and smooth-striated tubules, as previously alluded to (10,15,16) (Fig. 1c,d). Herein, proximal-striated tubule refers to those in proximity to the posterior salivary duct (PSD), and the distal-secretory tubule are distal to the PSD (Fig. 1b-e and Fig. S2). Interestingly the micro-CT reconstruction of the PSG in *E. berryi* allowed the identification of proximal-striated tubules but challenged the identification of secretory tubules, which are shown prominently in *E. berryi* tetrachrome staining (Fig. 1e and Fig. 2e,f). Additional hematoxylin and eosin staining revealed a consistent branching architecture of numerous small tubular structures (100 – 150 μm) with heterogenous diameters and lumen sizes in *O. bimaculoides* and *D. pealeii* PSG, illuminating the differentiation between the described tubule types (Fig. 2a-d). Tetrachrome staining of PSG sections in *E. berryi* further display granules, high protein content, and distinctive tissue layers between tubules potentially involving mucopolysaccharids (Fig. 2e,f). Our findings align with previous histological descriptions of the PSGs in the common cuttlefish (*S. officinalis*) (17) and the common octopus (*Octopus vulgaris*) (18). On a cellular level, transmission electron microscopy (TEM) images confirm an affluence of secretory granules and mitochondria in the secretory epithelial cells of *E. berryi, O. bimaculoides* and *A. bandense* (Fig. 2g-i). Taken together, the general morphology of venom glands across coleoid taxa is consistent in tubular distribution and structure, with few differences on a cellular-level such as abundance of secretory granular and mitochondria, which may vary based on different lineages, their stage of development, the venom release mechanism, the position within the gland or type of tubule.

**Figure 2.**
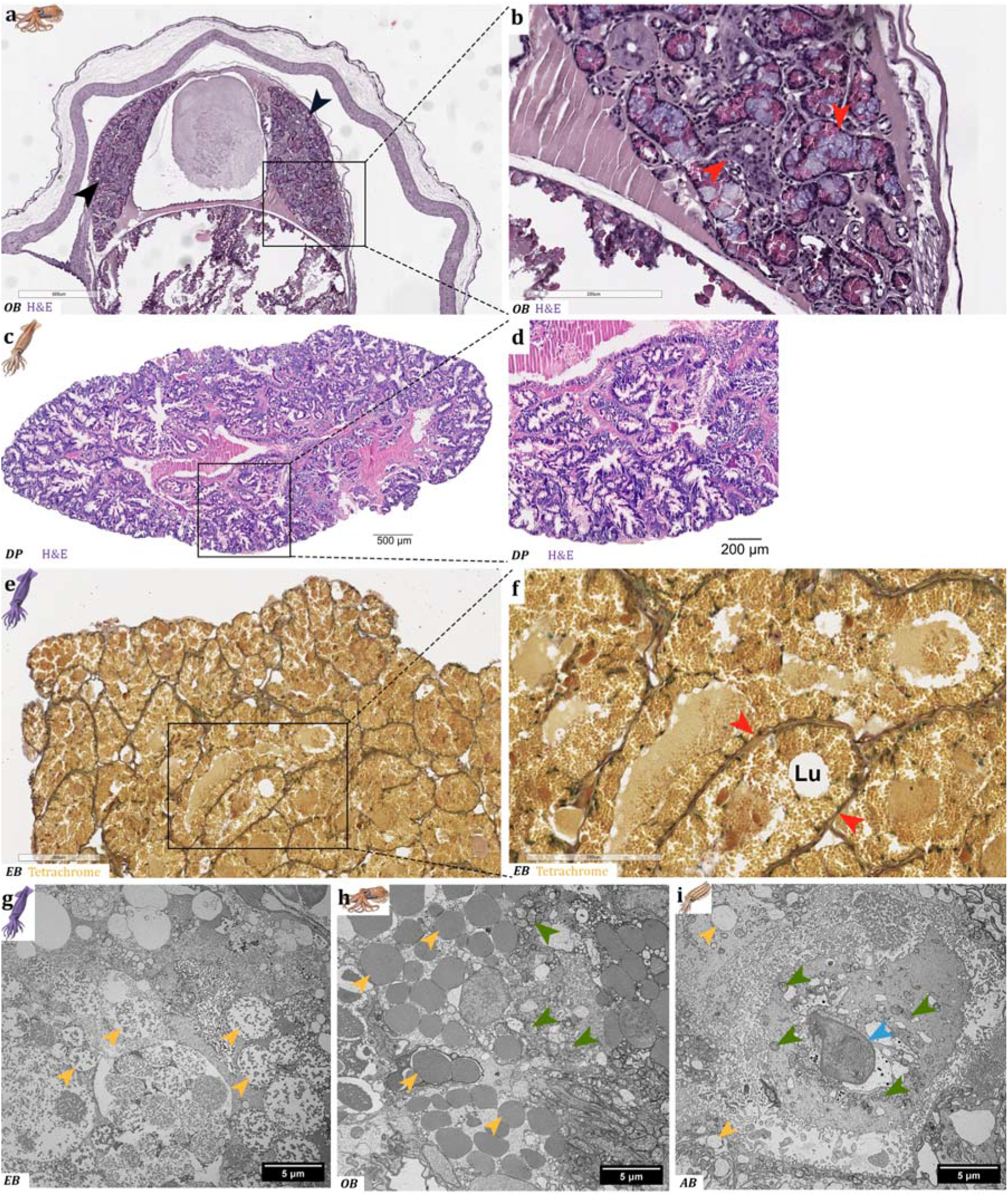
Histological characterization of coleoid posterior salivary glands (PSGs) reveal cross-lineage structural similarities. **a**,**b**, Hematoxylin and eosin staining of a formalin fixed, paraffin-embedded section of the PSG of adult *Octopus bimaculoides* and **c**,**d**, in *Doryteuthis pealeii*. Arrowheads: PSG (black), tubules (red). **e**,**f**, Tetrachrome staining in a tissue section of the PSG of an adult sepiolid squid *Euprymna berryi*. Lu = Lumen, arrowheads = tubules (red). **g-i**, Transmission electron microscopy images from the PSG of *E. berryi* (e), *O. bimaculoides* (f), and *Ascarosepion bandense* (g) highlighting abundant secretory granules and mitochondria. a-d, were obtained by high resolution digital slide scanning technique. Arrowheads: secretory granules (yellow), mitochondria (green), nucleus (blue).

### Neuro(muscular) structures and communication sites throughout coleoid venom glands

To determine if venom release is an actively regulated physiological process integrated with the nervous system, or a purely glandular phenomenon we characterized the muscular and/or neuromuscular structures in coleoid venom glands of *O. bimaculoides* and *D. pealeii* we used phalloidin and α-actin staining, respectively. Phalloidin highlights mostly the polymeric-filamentous state of actin in the venom gland, while the α-actin isoform is commonly associated with smooth muscle in vascular and visceral tissue. Our findings revealed an abundance of muscle fibers in *O. bimaculoides* that were specific and localized around tubules, matching DAPI-stained cell body aggregation (Fig. 3a). The α-actin staining in *D. pealeii* is strongly co-localized with the expression of co-stained venom protein SE-cephalotoxin-1 (*ctx-1*) and close to the basal membrane of gland tubular structure when contextualized with pkh26 membrane marker (Fig. 3b). Integration of the filamentous and α-actin stains with the spatial context of the gland structure provides molecular evidence for the presence of proposed circular smooth muscle layer around the secretory/striated tubules as observed morphologically in octopus PSG (10). Immunohistochemical staining with SMI-31 antibody targeting the heavy chains of neurofilaments, common structural elements found in axons and dendrites of nerves, molecularly validated the dense axonal infiltration previously described only in octopus (10,19) and now confirmed for the gland in *D. pealeii* (Fig. 3c). In addition, NeuN staining - marking mature neuron nuclei and perinuclear cytoplasm - in the PSG tissue sections of *A. bandense* is prominent as small, round clusters near the peripheral tubular openings and extends between the tubular structures until the center of the gland, with luminal areas devoid of staining (Fig. S3a). Similarly, in *D. pealeii* PSG, darkly stained round structures at the gland periphery suggest Nissl bodies – large aggregates of endoplasmic reticulum and ribosomes – are present in neuronal somas whose high rRNA content was targeted (Fig. S3b). While additional markers are needed to distinguish neuronal from secretory cell-related expression, these findings elucidate the presence of muscular and neuromuscular structures and neuronal communication sites in coleoid venom glands.

**Figure 3.**
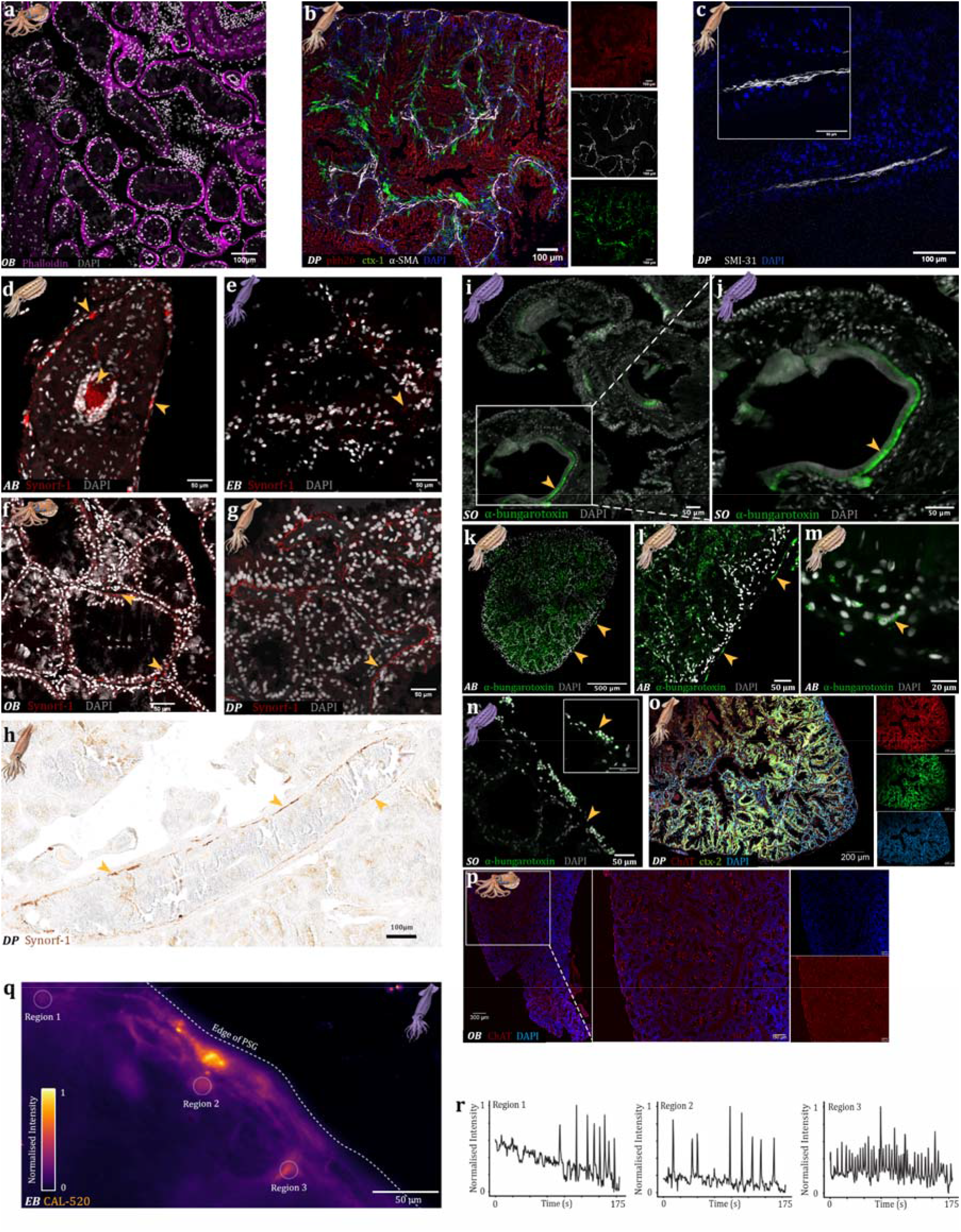
A network of neural agents is present in the posterior salivary glands (PSGs) of coleoid cephalopods. **a**, Muscle fiber distribution throughout the PSG in *Octopus bimaculoides* was visualized by staining actin filaments (phalloidin: purple) along with cell nuclei (DAPI: grey) for highlighting cell density distribution. **b**, Multiplexed immunohistochemistry and RNA-hybridization experiment in *Doryteuthis pealeii* PSG tissue section with the membrane cell linker pkh26 (red), alpha-actin smooth muscle marker (grey), SE-cephalotoxin-1 (*ctx-1*, green), the highly expressed venom protein in cephalopods and cell nuclei stained with DAPI. Single channel recordings are highlighted on the right side of this panel. **c**, Heavy chain of the neurofilament (grey) found in the PSG of *D. pealeii*. **d-h**, Synapses were visualized by staining phosphoproteins known to be abundant in presynaptic cytoplasm (synapsin) along with nuclear staining (DAPI: grey) in **d**, tentacle of adult *Ascarosepion bandense* (AB); and PSG of adult **e**, *Euprymna berryi* (EB), **f**, *O. bimaculoides* (OB), and **g, h**, *D. pealeii* (DP). **i-n**, Neuromuscular junctions visualized by labeling the alpha subunit of the muscle-type nicotinic acetylcholine receptor with α-bungarotoxin (α-nAChR: green) with cell density revealed by nuclear staining (DAPI: grey) in **i, j**, arm suckers and **n**, PSG of adult *Sepia officinalis* (SO), and **k-m**, *A. bandense* under different magnification. **o**, RNA-Hybridization chain reaction targeting choline acetyltransferase (ChAT: red) and SE-cephalotoxin-2 (*ctx-2*, green) in adult *D. pealeii* PSG and **p**, ChAT expression in adult *O. bimaculoides* at larger magnification (middle) from the overview image (left), and with the single DAPI-cell nuclei stained and ChAT channel recordings (right panels). **q**, Maximum intensity projection recorded in *ex vivo* PSGs of adult *E. berryi* during calcium imaging. **r**, Mean normalized intensity over time of regions 1-3 highlighted in p. Prominent signals highlighted with yellow arrowheads. The original calcium imaging recording in the ex vivo PSG can be found in supplementary material, *i*.*e*. Movie S1.

### Network of neuronal agents reveals potential operating routes from the nerves to venom release

To identify potential neuronal pathways in cephalopod venom release, we assessed the synaptic organization between neuronal and muscular tissue within the PSGs of squid (*E. berryi, D. pealeii*) and octopus (*O. bimaculoides*) by mapping the spatial distribution of presynaptic vesicle-associated phosphoproteins (synapsin), which participate in neurotransmitter release (20) (Fig. 3). Synapsin staining in *A. bandense* tentacles confirmed marker specificity and revealed strong central and peripheral signals consistent with known neuronal bundles and epidermal innervation (Fig. 3d). The synapsin signals detected in octopus and squid PSGs were widespread around the tubular structures, aligning with densely clustered cell nuclei with no luminal staining (Fig. 3e-h). Throughout the taxa examined, synapsin distribution is congruent with actin patterns, with sharpest, punctual staining in the tubular periphery of the venom gland in *O. bimaculoides* and *D. pealeii* (Fig. 3f-h).

To characterize the postsynaptic receptors, we examined muscle-type α-nicotinic acetylcholine receptors (nAChRs), which are abundant at neuromuscular junctions (NMJs), using the snake venom neurotoxin, α-bungarotoxin that binds with high affinity to muscle nAChRs and with low affinity to certain neuronal nAChRs (21). We confirmed staining specificity in *S. officinalis* arm suckers (Fig. 3i-j), observing sharp peripheral staining where NMJs are present. In the PSGs of cuttlefish (*S. officinalis* and *A. bandense*), we detected α-nAChRs in the gland periphery and diffuse staining between tubular structures (Fig. 3k-n). Contextualizing the patterns obtained from these single markers within the histological overview of the venom gland described above suggests that the smooth muscle tissue found around the tubular system in coleoid PSGs is abundantly targeted by neuronal tissue including cholinergic neurons.

Using hybridization chain reaction (HCR), we mapped the expression of choline acetyltransferase (ChAT), which is essential for acetylcholine production in cholinergic neurons and NMJs, in squid *D. pealeii* PSG. We found that the expression appears higher at the periphery of the PSG, and less intense throughout the gland (Fig. 3o). To determine if ChAT expression correlated with venom gene expression, probes for *D. pealeii* SE-cephalotoxin-2 (*ctx-2*) sequences were used to visualize this highly expressed venom transcript in the gland sections along with cell nuclei (DAPI). While ChAT expression supports the presence of acetylcholine-producing motor neurons, single-channel recordings showed limited differentiation between *ctx-2* and ChAT patterns, which may indicate proximity in the expression of venom protein *ctx-2* and cholinergic neurons (Fig. 3o). We repeated this experiment in *O. bimaculoides* PSG, with ChAT probe and DAPI only, as SE-cephalotoxin-like proteins have not been found in *O. bimaculoides* (22) (Fig. 3p). The results are comparable to the ones obtained for *D. pealeii*, but with lower intensity for ChAT expression. Taken together, these results provide robust evidence of a network of neuronal components that can act in concert to facilitate venom release from coleoid PSGs.

### Acetylcholine and dopamine-gated channels are present in coleoid venom glands

To identify neuronal ion channel family diversity in the PSGs of coleoid cephalopods, we retrieved cys-loop ligand-gated ion channels (cys-loop LGICs) sequences, including isoforms, from the PSG transcriptomes from *O. bimaculoides, D. pealeii*, and *A. bandense*, and performed a comparative phylogenetic analysis with other metazoan phyla. This allowed us to categorize the PSG sequences and predict their functional properties. The tree was split into three major families (Fig. 4): nicotinic ACh-like which are mainly cationic, GABA-A-like which are mainly anionic and cys-less which has an unknown function (23).

**Figure 4.**
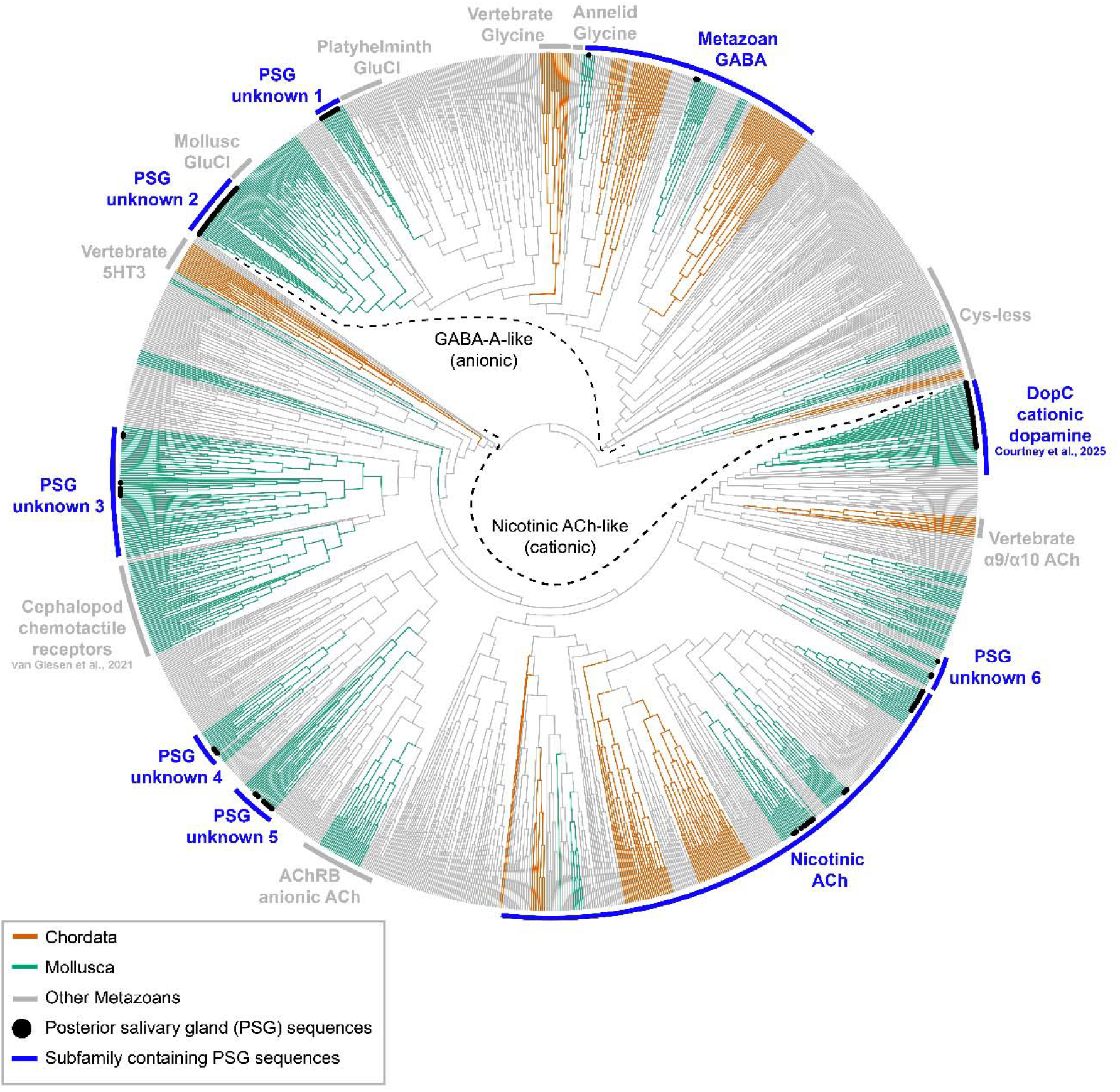
Comparative phylogenetics of cys-loop ligand gated ion channels from major metazoan phyla. Using venom gland (PSG) transcriptome sequences from *Octopus bimaculoides, Doryteuthis pealeii* and *Ascarosepion bandense* we identified excitatory acetylcholine and dopamine gated ion channels similar to those found in other metazoans. GABA-A-like channels associated with anionic activity are shown on the top half of the tree, while nACh-like channels associated with cationic activity are on the bottom. Further, several PSG sequence clusters were recovered with uncharacterized function but sister to either glutamate-gated channels (PSG unknown group 1 and 2), cephalopod chemotactile receptor group (PSG unknown 3) or inhibitory acetylcholine receptor group B (PSG unknown 4 and 5). Chordata sequences are highlighted in orange, molluscan sequences are in green, and all other metazoan sequences are in grey. PSG sequences are highlighted as black dots at the end of leaves, while subfamilies containing PSG sequences are highlighted in blue. PSG: posterior salivary gland.

GABA-A-like channels associated with anionic activity are shown on the top half of the tree (Fig 4). We identified a total of 46 sequences from which 3 sequences were found in the GABA-A group. We identified a cephalopod-specific subfamily (PSG unknown 1) with 11 PSG sequences from *A. bandense*, which is a sister to a glutamate-gated chloride channel group from platyhelminths (24). Interestingly, we also identified another cephalopod-specific group (PSG unknown 2), with 32 sequences from all three cephalopod species, which is sister to glutamate-gated chloride channels from molluscs (25). Further characterisation is needed to determine the functional properties of these anionic ‘unknown’ groups. Glycinergic cys-loop LGICs are present in vertebrates and annelids (26), but we didn’t see any evidence for these channels in cephalopods in this analysis.

Nicotinic acetylcholine-like channels associated with cationic activity are on the bottom of the tree (Fig. 4). We identified a total of 84 sequences from octopus, squid, and cuttlefish species, of which 26 sequences fall within the metazoan nicotinic acetylcholine-gated channel group. We also found in all three taxa of cephalopods tested, 34 sequences in the dopamine-gated channel group, which was recently functionally characterised in *O. vulgaris* and other invertebrates (27). There were also four groups with unknown functional properties (PSG unknown 3-6). The PSG unknown groups 4 and 5 were recovered as sister to the inhibitory acetylcholine receptor group B, while the PSG unknown group 6 is sister to a clade comprising the dopamine-gated channels and vertebrate α9/10 acetylcholine gated channels. The PSG unknown 3 group is of particular interest as it is recovered as sister to Cephalopod chemotactile receptors (28), which in the context of chemotactile stimuli from the arms could be involved in signal transduction for venom release in the PSG. Chemotactile-induced versus other forms of release of venom is an ongoing area of exploration in our group requiring further investigation. In contrast, the anionic acetylcholine receptor group previously found in molluscs (27,29) and the serotonin-gated channel group present in vertebrates and nematodes (30), were absent in PSG sequences and likely not present in cephalopod venom glands. These findings provide initial molecular and functional characterization of the ion channels found in squid, cuttlefish and octopus venom glands.

### Live staining of squid venom gland explant suggests calcium signaling activation

To monitor neuronal signal transmission, we examined if calcium signaling could be recorded in *ex vivo* cephalopod venom glands. Specifically, we incubated explanted PSGs from *E. berryi*, with a cell-permeable fluorogenic calcium indicator. Calcium plays a versatile role in the nervous system, but also in muscular tissue, acting as a second messenger and facilitating presynaptic exocytosis, postsynaptic signaling, and action potential development (31,32). We observed spontaneous calcium activity, particularly in areas surrounding nuclei near the edge of the PSG (Fig. 3q,r; see Movie S1). We also observed waves of calcium activity in thin processes travelling between these regions. Further analyses are required to determine if this activity is neural or muscular, and to identify the neurotransmitters or neuromodulators involved. While our *ex vivo* analysis is a pilot study, it indicates our ability to detect and measure signaling pathway activation and modulation in a real-time live staining of cephalopod venom gland explants, a previously unreported finding.

## Discussion

Coleoid cephalopods (octopuses, squids, and cuttlefish) use venoms to immobilize prey, however the mechanism(s) of venom secretion is unknown. Here, we combine multimodal histological profiling and gene expression analyses with calcium activity imaging and ion channel phylogenetics to elucidate venom gland architecture, innervation patterns, and neuronal signal transduction. Our data provides comprehensive experimental evidence for the hypothesis of neuromuscular control of cephalopod venom secretion morphologically assessed by Young over 60 years ago (7,9). Our results provide the first cross-lineage evidence that octopuses, squids, and cuttlefish share highly conserved microanatomical components of their venom gland (PSG) (Fig. 1 & 2). As a general pattern and in congruency with previous findings in *Octopus* spp. and *S. officinalis* (15,17), the glandular structure is strongly dominated by branching tubules lined with distinctive cell types that are packed with a high content of secretory granules. These tubules have varying dimensions and luminal area sizes that are potentially dependent on tissue section orientation and apical or basal localization with respect to the transition to the PSD. While the differentiation of proximal-striated and secretory tubules described in *O. vulgaris* (7,10) were identified in our *O. bimaculoides* histological analyses and micro-CT reconstructions, these features have yet to be fully verified within squid and cuttlefish PSGs. Tissue staining results in these lineages indicate a large proportion of secretory tubules but tubular differentiation within the micro-CT of *E. berryi* has been inconclusive. Further, the tandem usage of H&E and tetrachrome staining, and TEM for gland structure characterization highly support (chemical) content differentiation in the tubules (Fig. 2 & S2) but needs further optimization in the context of large PSG enzymatic content and species-specific tissue fixation success (see supplements, Fig. S4). However, the suggested differentiation observed in our findings alludes to distinct functions for parts of the tubular system with distal secretory tubules representing the production sites and the proximal smooth-striated tubule collect and propel the venom towards the salivary duct, underscoring a potential muscle-driven mechanism for venom release.

The infiltration of the venom gland by a connectivity tissue matrix enclosing blood vessels in proximity of muscle fibers targeted by nerve endings has been morphologically proposed and shown recently in single species of octopus and cuttlefish (15,17). However, a molecular verification across coleoid representatives has not been reported. Our muscular evidence identifies a high abundance of α-actin commonly found in smooth muscle fibers across the PSG of *D. pealeii* and the actin forms filamentous polymers in the PSG of *O. bimaculoides*. This matches the morphological observations from Young (7) and House (10) of circular layer of smooth muscle around the tubules in the PSG of *O. vulgaris*.

Another novel finding of muscular cholinergic receptors in the PSG of cuttlefish around the tubules match the patterns obtained from neuronal presynaptic vesicles and define clear neuromuscular communication sites in the proximity of the basal membrane of the gland tubular system. Together with the first indicators for presynaptic choline acetyltransferase signals in *D. pealeii* and *O. bimaculoides*, we present strong evidence that acetylcholine is a major neurotransmitter regulating the venom gland in cephalopods. This gains further support by our phylogenetic results indicating that fast excitatory cholinergic and dopaminergic signalling is likely playing an important role in the PSG, due to the presence of nicotinic acetylcholine and dopamine-gated channels. We also ruled out the possibility of fast cholinergic inhibition via AChRB channels, which were recently found to play a role in the *O. vulgaris* optic lobe (27). In addition, GABA and glutamatergic signalling may be acting as fast inhibitory signals, however more work is needed to confirm this. The other unknown groups represent interesting avenues for further exploration and may involve novel signalling mechanisms, especially in the context of chemotactile signal integration.

Current proposals for venom release include apocrine secretion, which involves budding of cellular cytoplasm, and is typical for mammalian salivary glands (33). However holocrine secretion, which involves bursting of cells, is proposed for cone snails (34). While the cellular secretion mechanism of venom in no venomous snail or cephalopod is known, the neural, neuromuscular and muscular structure we describe here support the hypothesis of neuromuscular control of the PSG in cephalopods, with muscle fibers around the secretory tubules and a marked innervation pattern across the gland. Our neuronal and neuromuscular recordings provide the first comparative insights into the interplay between neuronal and glandular systems in cephalopods, a significant first step in delineating the mechanisms of their brain-bite neural circuit for venom release. Similar observations of neural innervation in spiders (35), scorpions (36), snakes (37) and the giant centipede (38) venom glands point to a widespread phenomenon of neuronal control of venom release throughout vertebrates and invertebrates but with different complexity and molecular key players.

Together, these findings provide a multimodal foundation for interspecific comparisons of coleoid venom gland architecture, from the tubular differentiation to neuromuscular networks. By further resolving the molecular identity of smooth muscle fibers, axonal infiltration, cholinergic neuromuscular contacts, and potential calcium-dependent secretory pathways, our study substantially advances our understanding of how an external (chemotactile) stimuli may be integrated through the superior buccal lobe and translated into the activation of the secretory tubules. A cohesive working model for the transport of venom through the salivary ducts toward the buccal mass may occur by activation of the secretory tubules via cholinergic and dopaminergic channels, coupled with contraction of the surrounding smooth muscle.

Whether through pioneering work on the discovery of the squid giant axon by John Z. Young (39), or Alan Hodgkin and Andrew Huxley’s (40) use of the squid giant axon to characterize the mechanism of action potentials, the Cephalopoda nervous system has been used to advance groundbreaking knowledge. Our efforts are to bring to light the cephalopod neuronal and venom gland connection, which can be used to identify new circuits for animal predation, defense, and sexual competition. Our approach of integrating diverse datasets of histology, computed tomography models, phylogenetics and *ex vivo* calcium imaging harbors tremendous versatility for future studies in tracing neuronal routes from signal integration centers to specific venom gland structures, to characterize the molecular signaling networks and mechanistic sequence of cephalopod venom release. As such, these approaches deepen our understanding of neurosecretory systems in non-traditional model systems and hold considerable promises for expanding our understanding of cephalopod biology and accelerating biodiscovery within these evolutionary rich but underexplored lineages.

## Supporting information

Supplemental DataS1

LGC Full Phylogeny

LGC Alignment

## Resource availability

### Lead contact

Requests for further information and resources should be directed to and will be fulfilled by the lead contact, Mandë Holford.

### Materials availability

This study did not generate new, unique reagents.

### Data availability

- High-resolution digital scans of the *Octopus bimaculoides* hatchling and posterior salivary gland tissue sections are available at https://bio-atlas.psu.edu
- This paper does not report original code.
- Any additional information required to reanalyze the data reported in this paper is available from the lead contact upon request.

## Acknowledgments

The authors want to express their gratitude to the Marine Biological Laboratory’s Cephalopod Breeding, which resolved the husbandry of the specific species of squids (*E. berryi* and *D. pealeii*) cuttlefish (*A. bandense* and *S. officinalis*) and octopus (*O. bimaculoides*) used in this study and thus, represents a technological springboard for the detailed study of venom biology ex situ. M.H. also acknowledges MBL Whitman Fellow Kirsten Sadler Edepli for contributions to phalloidin staining. The authors would also like to thank Roger Hanlon and Stephen Senft for contributing the *O. bimaculoides* hatchling the Cheng lab team and 2-BM beamline scientists at Argonne National Laboratory for micro-CT based histotomography, Joshua Rosenthal for the access to the *E. berryi* micro-computed tomography and Jean Cooper for assistance with the digital scanning of presented tissue sections. The authors further acknowledge the Electron Microscopy & Histology services of the Weill Cornell Medicine Microscopy and Image Analysis Core, with special thanks to Juan Jimenez for his technical assistance, as well as the Harvard Center for Biological imaging and the PAIR-UP program at Marine Biological Laboratories allowing access to cutting-edge microscopy equipment and expertise. We would like to further thank Federica Pizzulli for the contribution of her expertise to fruitful and exciting discussions on the observed patterns.

## Author Contributions

Conceptualization, M.H. and K.N.K.; methodology, M.H., E.S., A.C., K.C., A.L.S., D.J.V., J.R.P.B., C.Ca., P. N., M.G., C.C. and M.W.; validation, M.H., E.S., A.C., K.C., A.L.S., D.J.V., J.R.P.B., C.Ca., C.C. and M.W.; formal analysis, K.N.K., M.H., E.S., A.C., K.C., A.L.S., D.J.V., J.R.P.B., C.Ca., C.C. and M.W.; investigation, K.N.K., M.H., E.S., A.C., K.C., A.L.S., D.J.V., J.R.P.B., C.Ca., P. N., M.G., C.C. and M.W.; resources, K.N.K. and M.H.; data curation, K.N.K, B.S., A.M., K.C., A.L.S., D.J.V., J.R.P.B., P. N., M.G., C.C. and M.W.; writing-original draft preparation, K.N.K, B.S., M.H. and M.W.; writing-review and editing, K.N.K., B.S., M.H., E.S., A.C., K.C., A.L.S., D.J.V., J.R.P.B., C.Ca., C.C. and M.W.; visualization, K.N.K., A.C., K.C., A.L.S., D.J.V., J.R.P.B., C.Ca., C.C. and B.S.; supervision, M.H.; project administration, M.H.; funding acquisition, M.H. All authors have read and agreed to the published version of the manuscript.

## Materials and Methods

### Ethical Statement

The animal study protocol was approved by the Institutional Review Board (or Ethics Committee) of The Marine Biological Laboratory (protocol code 2023_MBL-IBC-36 approved June 16, 2023).

### Animals and Tissue processing

Animal specimens were obtained from the Marine Biological Laboratory, Woods Hole (MA, USA). Euthanasia was performed by gradually increasing ethanol concentration from 0 to 30% in natural sea water over 30 min. If not otherwise stated in the following method protocols, for all tissue section staining, the PSGs from adult *D. pealeii, E. berryi, A. bandense, S. officinalis*, and *O. bimaculoides* were dissected and processed as described below. Briefly, after dissection, the gland samples were fixed overnight (at least 16 h) in 4% paraformaldehyde (PFA) in artificial salt water at 4°C. Fixed samples were washed with DEPC-treated PBS and gradually dehydrated through a progressive series of ethanol, of 10% increase per 20-min steps, up to 100%. To prepare the tissues for embedding, samples were cleared with two changes of histosol (30 min each), followed by incubation in two 1:1 histosol:paraffin solution changes (30 min each) and subsequent incubation in pure paraffin overnight at 60°C. The samples underwent four to five changes in 100% paraffin, each lasting 1 h, before being transferred into sectioning molds for solidification. Paraffin blocks were sectioned at 8-10 µm thickness using a microtome. For RNA-hybridization experiments, all water and phosphate buffered saline (PBS) solutions were treated with diethylpyrocarbonate (DEPC) overnight and autoclaved before use to eliminate RNase activity.

### (Immuno)histochemistry

Tissue staining procedures are described in brief below; detailed protocols are available in the Supplementary Material.

a. <u>Hematoxylin and Eosin staining</u>. For staining, tissue sections were deparaffinized in xylene rehydrated through a descending ethanol series and subsequently stained in Harris hematoxylin (Cat.-No. HHS32, Sigma-Aldrich, Missouri, USA) and counterstained with eosin Y solution (0.5–1%; Cat.No. 611815000, ThermoScientific, Waltham, USA). Tissue slides were mounted with DPX resin.
b. <u>Tetrachrome staining</u>. Tetrachrome staining procedure was performed according to the protocol established by Costa and Costa (41). This protocol involves sequential staining in four different dyes: Alcian Blue 8GX (Cat.-No. A5268, Sigma-Aldrich), Periodic acid (Cat.-No. 29922, Chem Impex), Schiff’s Reagent (Cat.-No. 3952016, Sigma Aldrich), Weigert’s Iron Hematoxylin (Cat.-No. 1.15973, Sigma Aldrich), and aqueous Picric acid (Cat.-No. 19552, Electron Microscope Sciences, Hatfield, USA). After concluding the staining procedure, slides were dehydrated with a progressive series of ethanol, cleared with xylene and mounted in DPX resin.
c. <u>Filamentous Actin</u>. Tissue sections were stained with phalloidin (Alexa Fluor 647, Cat -No. 8940, Cell Signaling Technology, MA, USA), counterstained with 4’,6-diamidino-2-phenylindole, dilactate (DAPI), mounted with hard-set mounting medium and coverslips. Fluorescence was recorded at 405 nm and 640 nm.
d. <u>Neurofilament-H</u>. Tissue sections were cleared with histosol and rehydrated through a descending ethanol series. Then, antigen retrieval and blocking were performed by using SMI-31 (Cat.-No. NE 1022, Sigma Aldrich) as primary and Donkey anti-Mouse IgG (H+L) Alexa 647 (Cat.-No. AP192SA6, Sigma Aldrich) as secondary antibody. Sections were mounted in DAPI-Fluoromount-G and coverslipped. Mounted slides were cured for approximately 24 h at 5°C before imaging. Fluorescence was recorded at 405 nm and 640 nm.
e. <u>Alpha-Pan Actin, Membrane and Cephalotoxin-1 multiplex experiment</u>. This involved immunohistochemistry (IHC), RNA-*in situ* hybridization chain reaction (HCR) and a membrane linker. Tissue sections were processed by a stepwise multiplexing strategy described in detail in the supplementary material. Briefly, first, the initial steps of the HCR protocol targeting cephalotoxin-1 were performed, beginning with deparaffinization, rehydration, and permeabilization (detailed below). To prevent heat-induced degradation or loss of integrity of the RNA probes used during HCR, the heat-based antigen retrieval of the IHC for the anti-pan actin antibody (Cat.-No. ab119952, Abcam, Waltham, USA) as primary antibody was carried out after these initial HCR steps. Following antigen retrieval, the HCR procedure was completed, including probe and amplifier hybridization. IHC staining was then concluded, starting from the blocking phase and followed by incubation with primary and using the Donkey anti-Mouse IgG (H+L) Alexa 647 (Cat.-No. AP192SA6, Sigma Aldrich) as secondary antibody. The multiplexing workflow was finalized through the addition of the pkh26 cell membrane labelling (Cat.-No. MIDI26, Sigma Aldrich), after which a nuclear counterstain was performed by mounting the tissue slides with DAPI-Fluoromount-G medium and finally slides were coverslipped. Fluorescence was recorded at 405 nm (DAPI), 488 nm (ctx-1), 561 nm (pkh26) and 640 nm (pan-actin).
f. <u>Synapsin</u>. Herein, we used an anti-SYNORF1 antibody (Cat.-No. 3C11, Developmental Studies Hybridoma Bank) that recognizes synapsin-1. This antibody is derived from the peptide sequence LFGGMEVCGL, located within the conserved C domain of synapsins. Its specificity has been validated across multiple species, including cephalopods. The secondary antibody was a goat anti-mouse IgG purchased from ThermoFisher (DyLightTM 650). The experimental method followed the antigen retrieval and blocking as described by Rees et al. (42), with few modifications to suit the objectives of this study. Finally, the slides were mounted in DAPI-Fluoromount-G, coverslipped, and stored at 5°C and left to dry for approximately 24 h before imaging. Fluorescence was recorded at 405 nm and 640 nm.
g. <u>Visualization of nAChRs at Neuromuscular Junctions.</u> We used α-bungarotoxin conjugates (Cat. No. B35450, ThermoFisher) to label nAChRs and visualize neuromuscular junctions. α-Bungarotoxin, a peptide derived from snake venom, binds irreversibly and with high specificity to the subunit alpha of the nAChRs in muscular tissue, but has also been shown to bind with lower affinity to nAChRs in the vertebrate brain (43), making it a valuable tool for their visualization in situ. The rehydration, antigen retrieval, and blocking stages followed the same protocol as of the synapsin. Fluorescence was recorded at 405 nm and 640 nm.
h. <u>NeuN</u>. Here, we used a Chromagenic DAB (ab64238; Abcam) staining to detect the presence of neuronal nuclei with the primary NeuN antibody (ABN91; Merck Millipore) by a standard IHC protocol including deparaffinization of tissue slides with xylene, antigen retrieval, and blocking phases. Donkey Anit-Chicken IgY (H&L) (Cat.No. 703-065-155, Jackson ImmunoResearch, West Grove, USa) was used as secondary antibody.
i. <u>Nissl bodies</u>. Squid PSG sections were used to perform a Cresyl Violet Stain (ab246817; Abcam). The slide first underwent deparaffinization and rehydration before proceeding with the Cresyl Violet Stain. The slide was then incubated in xylene before being cover slipped with CytoSeal XYL overnight.
j. <u>Choline Acetyltransferase and Cephalotoxin-2</u>. The method was initially developed by Choi et al. (44) with modifications according to Criswell and Gillis (45) and followed here accordingly. Hybridization chain reaction (HCR) reagents were obtained from Molecular Instruments Inc. (LA, USA), where probes for the venom protein cephalotoxin-2 for *D. pealeii* were further specifically designed. The probes for choline acetyltransferase were previously designed for *Octopus bocki* (46), and here tested on *D. pealeii* PSG sections. Sequences upon which the hybridized probes were designed can be found in the supplemental Data S1, while the comparative alignment of the *O. bocki* homolog ChAT sequences from *D. pealeii, O. bimaculoides and A. bandense* can be found in the supplementary methods and Fig. S5, while the ChAT cephalopod homolog sequences are found in Dataset S1. Fluorescence was recorded at 405 nm (DAPI), 488 nm (ctx-2) and 640 nm (ChAT).
k. <u>Live calcium signaling</u>. PSG was dissected from *E. berryi* specimen immediately after euthanasia and placed in a petri dish with a cephalopod-specific cell culture medium, herein called octomedia (see supplements for recipe). PSGs were incubated in the calcium indicator CAL-520 (Cat.-No. ab171868, Abcam, UK) that is known to improve signal-to-noise ratios compared to standard calcium imaging, allowing live fluorescence recordings of calcium migration within the PSG (47). Imaging was performed with an upright epifluorescent microscope equipped with a 490/525 nm excitation/emission filter set and operated at 2fps. Calcium imaging videos were processed as follows: image registration was performed within suite2p using the non-rigid registration approach (48), spontaneously active regions were manually defined, normalised using a min/max approach and plotted using a custom python script. The maximum intensity projection image shown in Fig. 3 was prepared for publication using ImageJ (49).

### Microscopy and Image Processing

Several different imaging techniques were employed to visualize the models, histological details, and staining results.

a. <u>Electron Transmission Microscopy</u>. Samples were washed with PBS, then fixed with a modified Karnovsky’s fix of 2.5% glutaraldehyde, 4% PFA and 0.02% picric acid in 0.1M sodium caocdylate buffer at pH 7.2 (50). Following a secondary fixation in 1% osmium tetroxide, 1.5% potassium ferricyanide, samples were dehydrated through a graded ethanol series and embedded in an epon analog resin. Ultrathin sections were cut using a Diatome diamond knife (Diatome, USA, Hatfield, PA) on a Leica EMEC7 ultramicrotome (Leica, Vienna, Austria). Sections were collected on copper grids and contrasted with lead citrate (51) and viewed on a JEM transmission 1400 electron microscope (JEOL, USA, Inc., Peabody, MA) operated at 100 kV. Images were recorded with a Veleta 2K x2K digital camera using RADIUS software (EMSIS, Germany).
b. <u>Micro-Computer Tomography.</u> An *O. bimaculoides* hatchling was obtained from the Marine Biological Laboratory and anesthetized in sea water with several percent ethanol and placed in a fixative of 4% paraformaldehyde in sea water for several days. The hatchling was immersed for a week in phosphotungstic acid and eosin solution, rinsed several times with distilled water, and then chemically dehydrated with acidified 2-2-di-methoxypropane (DMP) overnight. It was then immersed in hexamethyl di-silazane (HDMS) and air dried overnight in a hood. The intact animal was glued to an aluminium SEM stub mounted with plasticine onto a ThorLabs half inch diameter optical post screwed into a 1×1” kinematic base for imaging. Parallel beam synchrotron X-ray imaging was performed at sector 2 bending magnet beamline (2BM) of the Advanced Photon Source (APS) of Argonne National Laboratory. The beam energy was monochromatic and set to 25.5 keV and a sample to scintillator distance of 70 mm was used. Two sets of 9001 X-ray projections were collected over 360° of sample rotation to reconstruct the hatchling. 28% overlap between tomograms was chosen to minimize stitching artifact. Each projection was captured using a custom wide-field detector and camera with a 10mm field-of-view (FOV) and 0.7 μm^4^ voxel size. The 3D reconstructions were calculated using the gridrec algorithm from TomoPy (52) and stitched with manual alignment in ImageJ. The Bronnikov Aided Correction was used for phase retrieval with alpha and beta parameters equal to 2.0 (53,54). A formalin-fixed, paraffin-embedded *E. berryi* sepiolid squid was obtained from the Rosenthal Lab at the Marine Biological laboratory. Excess paraffin was removed via scalpel and excess paraffin wax was used to affix the sample to a plastic sample holder. Parallel beam synchrotron X-ray imaging was performed at beamline 8.3.2 of the Advanced Light Source (ALS) of Lawrence Berkeley National Laboratory. The beam energy was monochromatic and set to 14 keV, and a sample to scintillator distance of 50 mm was used. 1969 projections with 300 ms of exposure time were captured over 180 degrees of rotation using a 5 mm FOV detector with 0.5 µm isotropic voxel size. The 3D reconstructions were calculated using the gridrec algorithm from TomoPy and stitched with manual alignment in ImageJ. The Bronnikov Aided Correction was used for phase retrieval with alpha and beta parameters equal to 2.0 and 1.25 respectively (53,54).
c. <u>Confocal microscopy</u>. The imaging of the phalloidin, SMI-31, multiplexed experiment, synorf-1 (Fig. 3d-g), NMJ and ChAT were performed using a Zeiss LSM 880 confocal microscope equipped with 10x, 25x, 40x, and 63x objectives. Image processing was performed with the Zeiss Zen 3.10 blue version and ImageJ software.
d. <u>Light microscopy</u>. Synorf-1 staining in *D. pealeii* (Fig. 3h) was captured with a Zeiss Axioscope 5 at a resolution of 1024 × 1024 pixels. Images were processed with Image J Fiji software.
e. <u>Digital Slide scan</u>. Tetrachrome and H&E tissue slides were scanned at 40× using an Aperio AT2 slide scanner (Leica Biosystems, Nussloch, Germany) and images were saved in TIFF format. Image processing was made with the manufacturer ImageScope software. NeuN and Nissl body stainings were visualized on a digital pathology scanner (NanoZoomer-SQ Digital slide scanner, Hamamatsu Photonics K.K., Shizuoka, Japan).

### Comparative phylogenetic analysis of metazoan cys-loop LGICs

We used the ‘cys-loop’ LGIC dataset from Courtney *et al*. (27) which included 26 species of metazoans across all major phyla and 1197 sequences. We added cys-loop LGIC sequences identified from *O. bimaculoides, D. pealeii*, and *A. bandense* PSG transcriptomes to this dataset (see supplements for methodological details). Alignment, trimming, tree generation and visualization were performed as described in Courtney *et al*. (27). In brief, alignment was performed using MAFFT (v7.505) with E-INS-i parameters (54), trimming was performed using Trimal (v1.4.1) gappyout method (56). Phylogenies were generated using IQ-TREE v2.1.4_beta with 1,000 ultra-fast bootstraps (57). The tree model was calculated using IQ-TREE’s ModelFinder implementation according to Bayesian inference criterion, and LG+F+R10 was used. The tree was visualized using TreeViewer (58). We used the cys-less group identified by Jaiteh *et al. (*23) to root the tree. The alignment of the metazoan cys-loop LGIC sequences and the phylogeny with full annotations are available as supplemental files.

