## Supplementary figures and images for "Neurons, Muscles, and Venom: Elucidating a Neural-to-Secretory Pathway in Cephalopod Predation"

### LGC Full Phylogeny

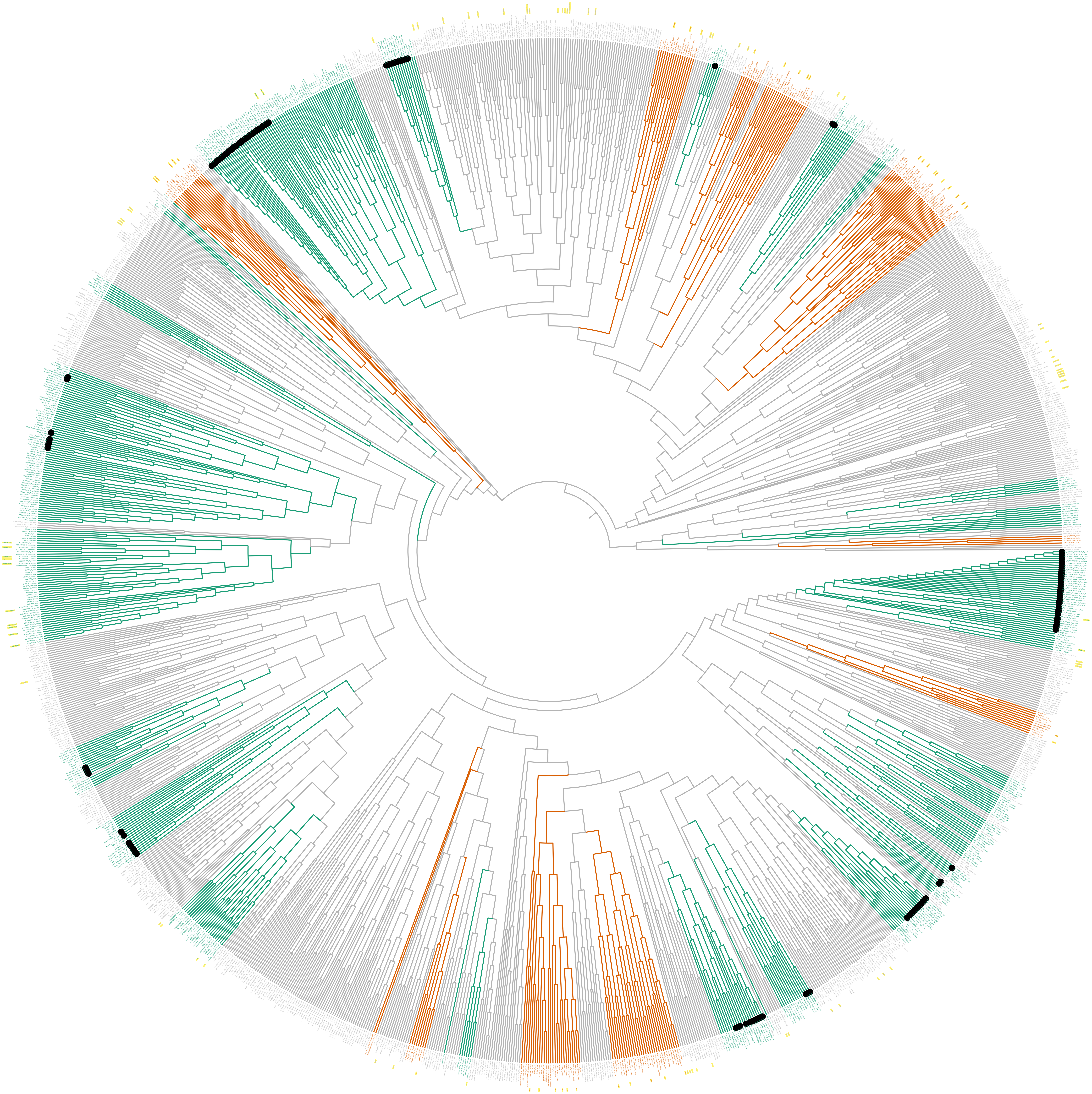
