## Supplemental Information and Methods for "Neurons, Muscles, and Venom: Elucidating a Neural-to-Secretory Pathway in Cephalopod Predation"

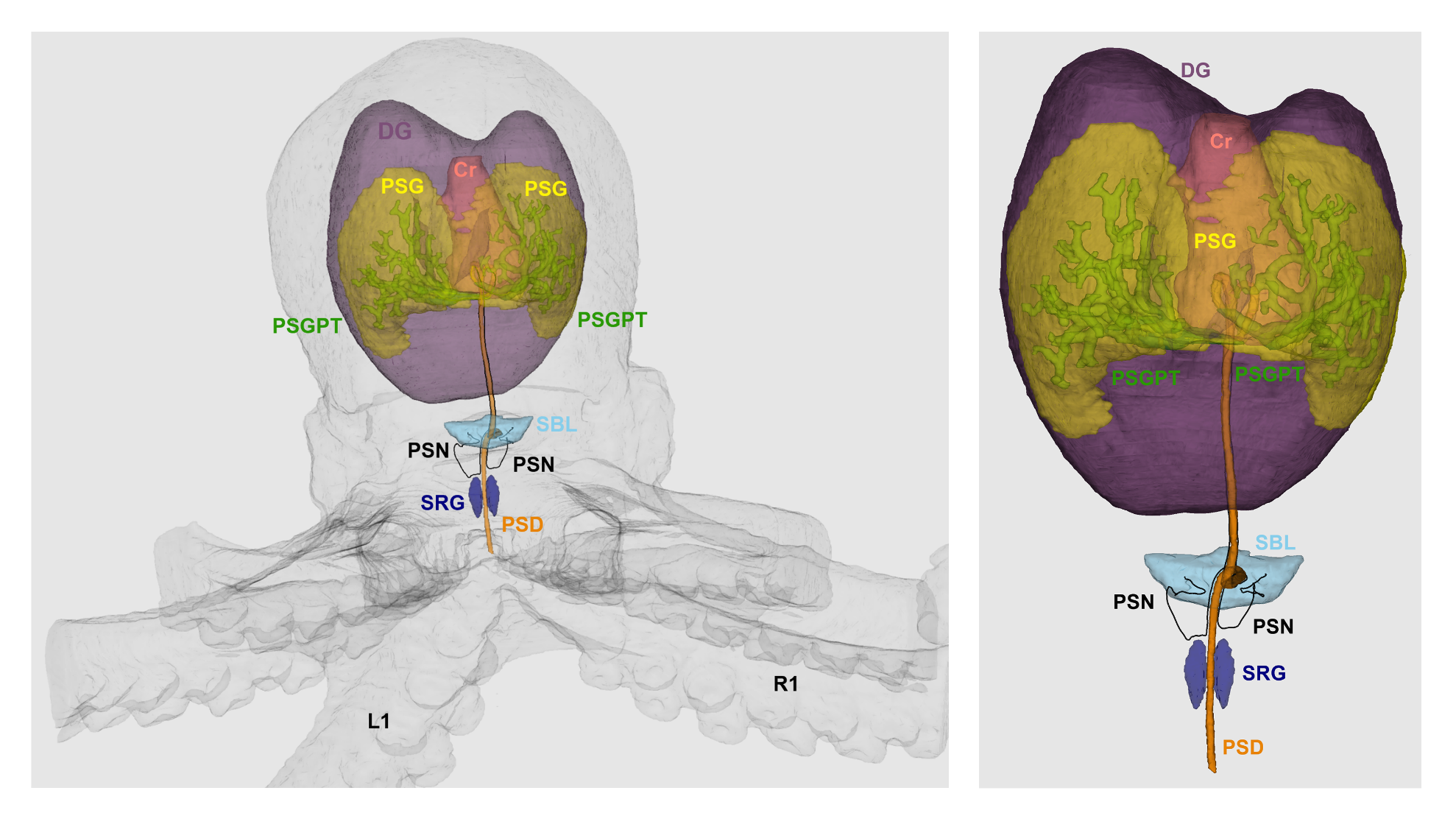

**Figure S1. Venom release from the posterior salivary gland in cephalopods is potentially neuronally controlled.** Micro-computed tomography of an *Octopus bimaculoides* hatching (left) highlights the macroanatomical position of the posterior salivary gland (PSG), *i.e.* venom gland, the tubular system within the gland (PSGT), the salivary duct (PSD), as well as the potential neuronal control centers, *i.e.* superior buccal lobe (SBL) and posterior salivary nerves (PSN), and the subradular ganglion (SRG). For anatomical contextualization, we further highlighted the digestive gland (DG) and crop (Cr), and added a magnification of the *ex situ* venom system (right).

**Table S1.**

Overview of experiment coverage across cephalopod species within this study (green field denote the included analyses in the present study).

|  | **Myopsida**  **[Squid]** | **Sepiolida**  **[Bobtail squids]** | **Octopoda**  **[Octopus]** | **Sepiida [Cuttlefish]** | |
| --- | --- | --- | --- | --- | --- |
|  | ***Doryteuthis pealeii*** | ***Euprymna berryi*** | ***Octopus bimaculoides*** | ***Sepia bandensis  (Ascarosepion bandense)*** | ***Sepia officinalis*** |
| **H&E** |  |  |  |  |  |
| **Tetrachrome** |  |  |  |  |  |
| **TEM** |  |  |  |  |  |
| **Micro-CT** |  |  |  |  |  |
| **Synapsin** |  |  |  |  |  |
| **m-nAChr** |  |  |  |  |  |
| **F-Actin** |  |  |  |  |  |
| **Multiplex  (α-Pan Actin, pkh26, ctx, DAPI)** |  |  |  |  |  |
| **Neurofilament-H (SMI-31)** |  |  |  |  |  |
| **ChAT** |  |  |  |  |  |
| **Calcium signaling**  **(*ex vivo*)** |  |  |  |  |  |
| **NeuN** |  |  |  |  |  |
| **Nissl bodies** |  |  |  |  |  |
| **Phylogeny  cys-loop LGC** |  |  |  |  |  |
| **Tissue fixation** |  |  |  |  |  |

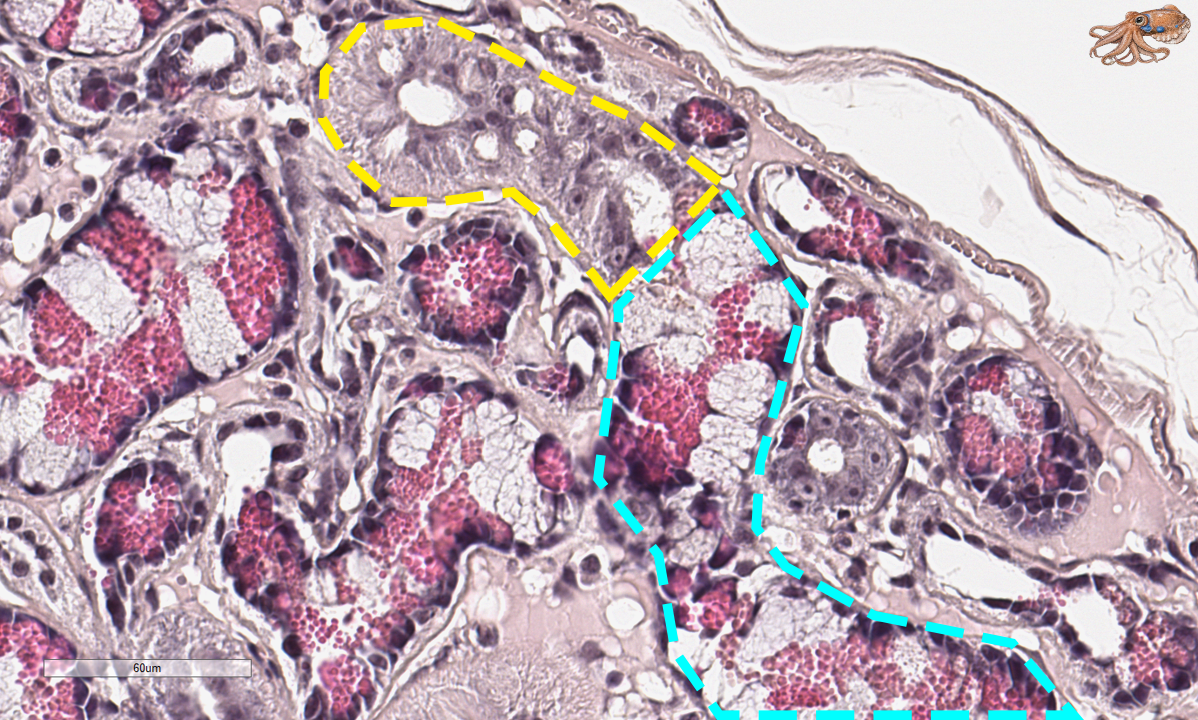

**Figure S2. Cephalopod posterior salivary glands exhibit two differentiate tubule types.** Hematoxylin and eosin staining of a formalin fixed, paraffin embedded posterior salivary gland tissue section of an *Octopus bimaculoides* hatchling highlighting with the yellow dashed line the transition between the two known tubule types: striated (yellow dash lined) and secretory (blue dash lined).

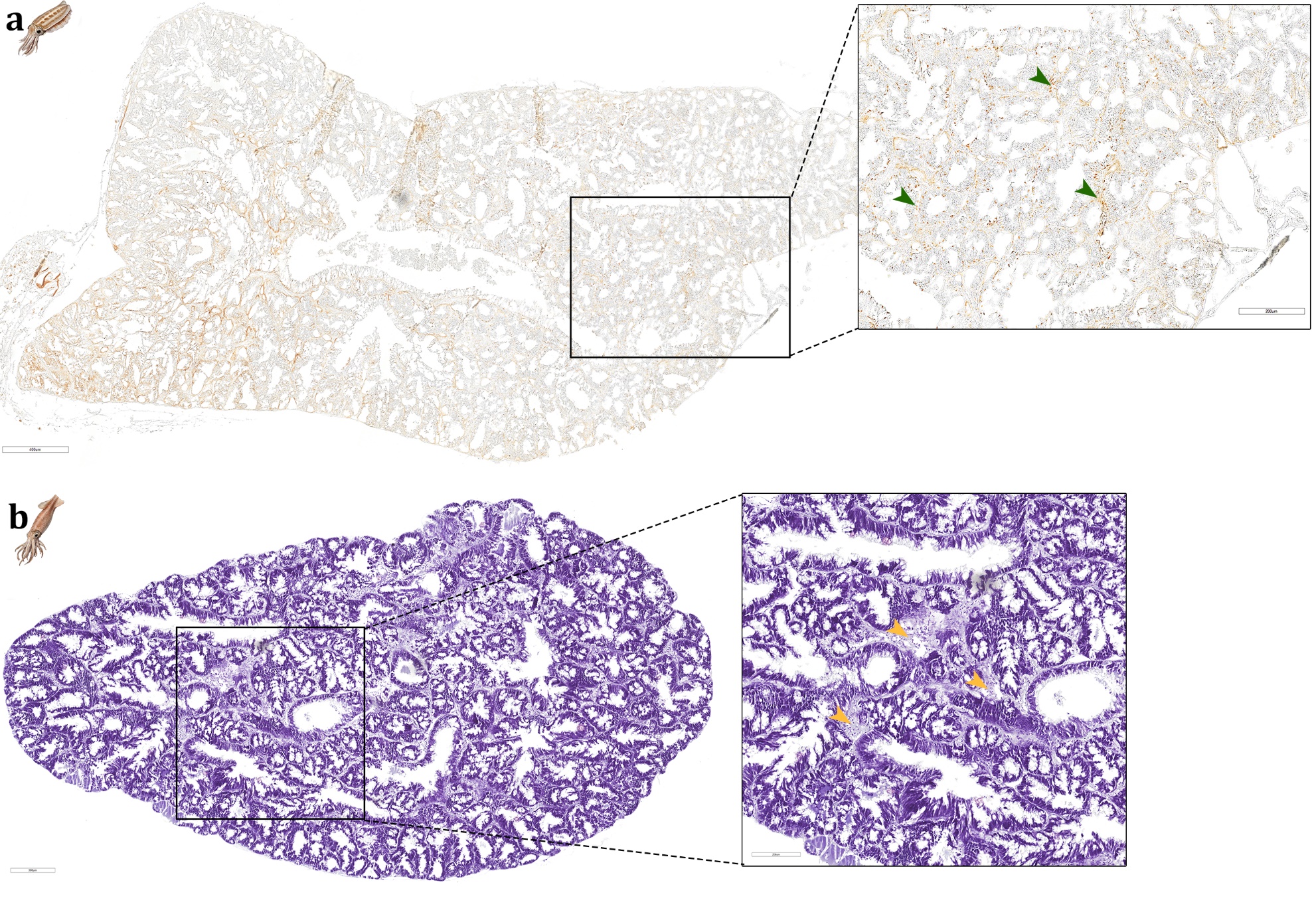

**Figure S3. Cephalopod posterior salivary gland paraffin sections targeted by neuronal markers: a,** Neuronal nuclei antigen for RNA-binding proteins immunoreaction (NeuN; brown) in adult *Ascarosepion bandensis*; **b,** Nissl bodies, large aggregates of rough endoplasmic reticulum and ribosomes common in neuronal cytoplasma (violet) and cell nuclei (dark purple) staining in adult *Doryteuthis pealeii*. Yellow and green arrowheads highlight distinctive signal areas.

***Supplementary methods***

***(Immuno)histochemistry***

a. Hematoxylin and Eosin staining. For staining, tissue sections were deparaffinized in xylene (2 × 5 min), rehydrated through a descending ethanol series (100%, 95%, 70%; 2 min each), and rinsed in distilled water. Slides were stained in Harris hematoxylin (Cat.-No. HHS32, Sigma-Aldrich, Missouri, USA) for 5–7 min, rinsed in running tap water for 5 min to stain the nuclei, and briefly dipped in 0.3% acidified alcohol to remove excess of dye. After another rinse with tap water, slides are being dipped into lithium carbonate for 1 min to increase staining intensity of Hematoxylin. Following another rinse in distilled water, slides were counterstained with eosin Y solution (0.5–1%; Cat.No. 611815000, ThermoScientific, Waltham, USA) for 30 s to 2 min. After staining, sections were sequentially dehydrated through an ascending ethanol series (30%, 95%, 100%; 1 min each), cleared in xylene for 1 min, and mounted with DPX resin.
b. Tetrachrome staining. Tetrachrome staining procedure was performed according to the protocol established by Costa and Costa (1). Briefly, tissue sections were sequentially introduced into: Alcian Blue 8GX (30 min; Cat.-No. A5268, Sigma-Aldrich), Milli-Q wash (2 x 30 s), 0.5% Periodic acid (5 min; Cat.-No. 29922, Chem Impex), Milli-Q wash (at least 5 immersions), Schiff’s Reagent (15 min; Cat.-No. 3952016, Sigma Aldrich), Milli-Q wash (2 x 30 s), Weigert’s Iron Hematoxylin (2 min; Cat.-No. 1.15973, Sigma Aldrich), running tap water (1 min), aqueous Picric acid (5 min; Cat.-No. 19552, Electron Microscope Sciences, Hatfield, USA), Milli-Q wash (at least five immersions). All dyes were applied in the dark and at room temperature (RT), with the exception of Picric Acid solution, which was preheated to 60°C before use to enhance staining. Alcian Blue was prepared with 1% m/v in 3% v/v acetic acid (the solution was filtered after preparation). The hematoxylin solution was prepared using a 1:1 ratio of Solution A and Solution B, as directed by Sigma. After concluding the staining procedure, slides were dehydrated with a progressive series of ethanol, cleared with xylene and mounted in DPX resin.
c. Filamentous Actin. For phalloidin staining (Alexa Fluor 647, Cat.-No. 8940, Cell Signaling Technology, MA, USA), tissue section of the PSG was incubated for 2 h at RT in the dark. Cells were counter-stained with 4’,6-diamidino-2-phenylindole, dilactate (DAPI) for 10 min at RT. After each step, the cells were washed three times with PBS and mounted with hard-set mounting medium and coverslips (1.5 mm). Fluorescence was recorded at 405 nm and 647 nm.
d. Neurofilament-H. Tissue sections were rehydrated through a descending ethanol series. Slides were immersed sequentially in Histosol (2 × 5 min), 100% ethanol (2 × 5 min), 75%, 50%, and 25% ethanol in PBS (5 min each), followed by two washes in Milli-Q water (2 min each). Sections were then rinsed in PBS containing 0.1% Triton X-100 (3 × 5 min).

Antigen retrieval was performed by incubating slides for 5 min in pre-heated distilled water at 60 °C, followed by transfer into pre-warmed sodium citrate buffer (10 mM, pH 6.0). Slides were incubated at 95°C for 25 min and subsequently cooled in a −20°C freezer for up to 10 min. After cooling, sections were rinsed in PBS with 0.1% Triton X-100 (2 × 5 min). For blocking, slides were placed horizontally in a humidified chamber and incubated for 1 h at RT with blocking solution (5% milk dissolved in PBS containing 0.1% Triton X-100). SMI-31 (Cat.-No. NE 1022, Sigma Aldrich) were diluted in blocking solution at 1:500, and 150–200 µL of solution was applied per section. Slides were coverslipped with parafilm and incubated overnight at 4°C in a humidified chamber. Following removal of primary antibody, sections were washed in PBS + 0.1% Triton X-100 (3 × 10 min). Donkey anti-Mouse IgG (H+L) Alexa 647 (Cat.-No. AP192SA6, Sigma Aldrich) was used as secondary antibody and diluted again at 1:500 in blocking solution were applied (150–200 µL per slide), slides were coverslipped with parafilm, and incubated for 4 h RT in the dark. Finally, slides were washed again in PBS + 0.1% Triton X-100 (3 × 10 min), followed by three extended washes of 30–60 min each to ensure complete removal of unbound secondary antibodies. Sections were mounted in Fluoromount-G (with DAPI) and coverslipped. Mounted slides were cured for approximately 24 h at 5°C before imaging.

e. Alpha-Pan Actin, Membrane and Cephalotoxin-1 multiplex experiment. This involved immunohistochemistry (IHC), RNA-*in situ* hybridization chain reaction (HCR) and a membrane linker. Tissue sections were processed by a stepwise multiplexing strategy described in the following. First, the initial steps of the HCR protocol targeting cephalotoxin-1 were performed, beginning with deparaffinization, rehydration, and permeabilization (detailed below). To prevent heat-induced degradation or loss of integrity of the RNA probes used during HCR, the heat-based antigen retrieval of the IHC for the alpha-pan actin antibody was carried out after these initial HCR steps. Following antigen retrieval, the HCR procedure was completed, including probe and amplifier hybridization. IHC staining was then concluded, starting from the blocking phase and followed by incubation with primary and secondary antibodies. The multiplexing workflow was finalized through the addition of the cell-linker labeling protocol, after which a nuclear counterstain was applied by mounting the tissue slides with DAPI-containing medium.

In more detail, the HCR was performed as following described: Slides were deparaffinized twice in histosol for 5 min. Slides were then rehydrated in a series of decreasing ethanol solutions: 100% ethanol for 5 min ×2, then 2 min each in 90%, 70%, and 50% ethanol/DEPC-PBS, DEPC water, and DEPC-PBS with 0.1% Tween20. Subsequently, slides were treated with 10 µg/ml proteinase K (25530049; Thermo Fisher) in DEPC PBS at 37°C for 10 min and rinsed in DEPC water at RT for 2 min. Now, antigen retrieval was performed as described for the Neurofilament staining (point d) with stepwise temperature transition from 60°C to 95°C and back to -20°C. Back in the next step of the HCR protocol, slides were moved to a humidified chamber for and pre-hybridized by adding 2 ml of pre-warmed hybridization buffer per slide. Slides were incubated in the humidified chamber at 37°C for 30 min. During this pre-hybridization step, probe sets were diluted in pre-warmed hybridization buffer, with 0.8 µl of each 1 µM probe stock added per 100 µl of hybridization buffer, preparing 300 µl of hybridization buffer per slide, and the mixture was returned to 37°C. After the 30 min pre-hybridization, the humidified chamber was removed from the 37°C incubator and 250 µl of hybridization buffer with the probe was added to each slide. A parafilm coverslip was set over each slide and they were incubated in a humidified chamber at 37°C overnight.  On the second day, parafilm was carefully removed, and slides were briefly washed twice with around 1 ml of pre-warmed HCR wash buffer. Then, slides were washed at 37°C with the pre-warmed solutions as follows: 75% wash buffer for 15 min, 50% wash buffer for 15 min, 25% wash buffer for 15 min, and 5x SSCT for 15 min. Slides were then washed in 5x SSCT at RT for 5 min. To prepare the hairpins, 8 µl of each hairpin solution (H1 and H2) were heated separately in microcentrifuge tubes to 95°C for 90 s and then cooled in the dark at RT for 30 min. Hairpin solution were proportionally prepared for a final volume of 200 µl of amplification buffer with hairpins per slide. Slides were moved to a humidified chamber and pre-amplified by applying 2.5 ml of RT amplification buffer to each slide. Slides were incubated in the humidified chamber for 30 min at RT. Hairpins and amplification buffer were then mixed and briefly vortexed, the pre-amplification buffer was drained off, and 200 µl of this mixture was added to each slide. A parafilm coverslip was added, and the slides were incubated in a humidified chamber overnight in the dark at RT. On the third day, slides were washed in the dark as follows: 5x SSCT for 5 min, twice with 5x SSCT for 15 min, and 5x SSCT for 5 min.

At this point, we returned to the blocking step in the antibody protocol as described above and concluded the subsequent IHC steps by using the anti-pan actin antibody (Cat.-No. ab119952, Abcam, Waltham, USA) as primary antibody and ). Donkey anti-Mouse IgG (H+L) Alexa 647 (Cat.-No. AP192SA6, Sigma Aldrich) as secondary antibody. The pkh26 cell membrane labelling (Cat.-No. MIDI26, Sigma Aldrich) was performed following the manufacturer’s protocol. Finally, glass coverslips (1.5 mm thickness) were applied with Fluoromount-G with DAPI. Slides were stored in the dark and allowed to dry overnight before being stored at 4°C in the fridge.

f. Synapsin. Herein, we used an anti-SYNORF1 antibody (Cat.-No. 3C11, Developmental Studies Hybridoma Bank) that recognizes synapsin-1. This antibody is derived from the peptide sequence LFGGMEVCGL, located within the conserved C domain of synapsins. Its specificity has been validated across multiple species, including cephalopods. The secondary antibody was a goat anti-mouse IgG purchased from ThermoFisher (DyLightTM 650). The experimental method was adapted from Rees et al. (2), with modifications to suit the objectives of this study.

The slides were rehydrated following a stepwise protocol of 2x histosol (5 min each), 2x 100% EtOH (5 min each), 75% EtOH/PBS (5 min), 50% EtOH/PBS (5 min), 25% EtOH/PBS (5 min), 2x washes in distillated water (2 min) and 3x washes in PBS supplemented with 0.1% Triton-X (5 min each). After preheating the tubes, the slides containing the PSG tissue sections were incubated in a 60°C distilled water bath for 5 min. The warmed slides were then transferred to pre-warmed antigen retrieval solution (10 mM sodium citrate, pH 6.0) and incubated at 95°C for 25 min. Subsequently, the slide container was cooled in a -20°C freezer for no more than 10 min, ensuring the slides did not freeze. The slides were then transferred to a glass Coplin jar and rinsed twice for 5 min each in PBS containing 0.1% Triton-X.

Next, the slides were placed in a humidified chamber, and 2 ml of blocking solution (10% sheep or goat serum in PBS with 0.1% Triton) was added to each slide, followed by incubation for 1 h at RT. For the primary anti-body application, slides were treated with the Synorf antibody diluted in blocking solution (9.75 µg/ml, 150-200 µl per slide). Each slide was covered with parafilm and incubated overnight at 4°C in a humidified chamber.

The following day, the slides were rinsed three times for 10 min each in PBS with 0.1% Triton, and secondary antibodies were prepared. The secondary antibody was diluted in blocking solution (3.3 mg/ml, 150-200 µl per slide), shortly centrifuged and applied to the slides, which were then covered with parafilm and incubated at RT in the dark for 4 h. Following this, the slides were rinsed three times for 10 min in PBS with 0.1% Triton-X, followed by three additional washes of 30-60 min each to ensure thorough removal of the secondary antibody. To ensure the experiment was working and detecting synapsin, we used brain sections as a positive control. For the negative control, additional PSG gland tissue sections from the same specimens were processed without applying the primary antibody to verify the specificity of the staining. Finally, the slides were mounted in Fluoromount-G (with or without DAPI), and glass coverslips (1.5 mm thickness) were applied. The slides were stored at 5°C and left to dry for approximately 24 h before imaging.

g. Visualization of nAChRs at Neuromuscular Junctions. In this study, we used α-bungarotoxin conjugates (Cat. No. B35450, ThermoFisher) to label nAChRs and visualize neuromuscular junctions in the PSG of *S. officinalis*, *A. bandense*, and *O. bimaculoides*. α-Bungarotoxin, a peptide derived from snake venom, binds irreversibly and with high specificity to the subunit alpha of the nAChRs in muscular tissue, but has also been shown to bind with lower affinity to nAChRs in the vertebrate brain (3), making it a valuable tool for their visualization in situ.

The rehydration, antigen retrieval, and blocking stages followed the same protocol as of the synapsin. A concentration of 1 µg/ml α-bungarotoxin staining solution in PBS was prepared, and approximately 150-200 µL were applied per slide. Arm tissue sections through the suckers were used as a positive control to confirm the detection of nAChRs, while PSG tissue sections processed without α-bungarotoxin served as negative controls to assess staining specificity.

The sections were covered with parafilm and incubated at RT in the dark for 4 h. After incubation, the sections were rinsed three times in PBS, mounted in Fluoromount-G, and covered with a coverslip.

h. NeuN. Here, we used a Chromagenic DAB staining to detect the presence of neuronal nuclei with a NeuN antibody (ABN91; Merck Millipore). Stock solutions for 0.3% Triton-X/PBS (X100; Sigma Aldrich) and 0.3% Tween20/PBS (P7949; Sigma Aldrich) were made one day prior to the start of the experiment. Calculations for 3% hydrogen peroxide (H325-100; ThermoFisher), blocking solution, NeuN primary antibody solution, and secondary antibody solution were conducted the day before. The next two days follow a standard immunohistochemical procedure:

On the first day of the experiment, the slide underwent a series of rinses to deparaffinize and rehydrate with Xylene, Ethanol, and Milli-Q Water. During these rinses, 1X PBS (P3813; Sigma Aldrich) and the antigen retrieval, which was diluted with 1X Citrate Retrieval Buffer (C9999; Sigma Aldrich), were prepared and placed inside a steamer (B0BSHJ5LQ5; Elite Gourmet) for 45 min alongside an additional container of Milli-Q water. Temperature probes were used to monitor the temperature to ensure it reached 90˚C (roughly 45 min before rehydration ended). The slide rack was placed in slide well with Milli-Q water inside steamer for 10 s before transferring to slide well with antigen retrieval for 10-15 min. During this time, a humidity chamber (Stain Tray Slide Staining System, Thermo Fisher) and 3% Hydrogen Peroxide were prepared. The slides were then removed from the antigen retrieval and cooled for 20 min before being placed in a slide well with 1X PBS for 5 min. The slide was then placed inside the humidity chamber and 3% Hydrogen peroxide was added for 10 min to remove the endogenous peroxide on the tissue. Once the slide was placed in a fresh slide well with 1X PBS for 5 min, the blocking Solution was prepared. After, the slide was moved to the humidity chamber and the blocking solution was placed on the slide. While the slide was incubating at RT for 1 h, the NeuN primary antibody (1:100 dilution) was prepared. After the incubation, the slide was moved to the humidity chamber and the NeuN primary antibody was added before being cover slipped and left inside the humidity chamber overnight at 4˚C. On the second day, the secondary antibody*, i.e.* Donkey Anit-Chicken IgY (H&L) (Cat.No. 703-065-155, Jackson ImmunoResearch, West Grove, USA) was prepared at a 1:500 dilution while the slide underwent 5 separate rinses with 1X PBS for 5 min each to wash off the primary solution. The secondary antibody was then placed on the slide and incubation was for 1 h before the slide was washed three times with 1X PBS for 5 min each. Then, 2-3 drops of Streptavidin (SA-5704; Vector Laboratories) were placed on the slide inside the humidity chamber for 30 min while Coplin jars were prepped with 1X PBS. As the slide was washed again in a similar sequence of 1X PBS rinses, the DAB solution (ab64238; Abcam) (40 µl chromagen to 40-45 drops of DAB substrate) was prepared. For the slide, it was placed under a microscope as the DAB solution was placed on top and sat for 1-2 min to allow for development and appearance of neuronal nuclei to appear (brown-orange color). Once complete, the slide was rinsed several times with 1X PBS before being placed in an additional slide well with 1X PBs before beginning rehydration. The slide then underwent a series of rinses with xylene and ethanol to rehydrate the slide. After, the slide was then prepared with CytoSeal XYL (18009; Electron Microscopy Sciences) and cover slipped overnight. The slide was then scanned on a digital pathology scanner (NanoZoomer-SQ Digital slide scanner, Hamamatsu Photonics K.K., Shizuoka, Japan) to observe the results.

i. Nissl bodies. Squid PSG sections were used to perform a Cresyl Violet Stain (ab246817; Abcam). The slide first underwent a series of rinses that were 5 min each to deparaffinize and rehydrate the slide using xylene and ethanol. Before proceeding with the Cresyl Violet Stain, the slide sat in a slide well with Milli-Q water for 5 min. After, the slide incubated in a coplin jar with 1% Cresyl Violet Stain solution for 20 min to ensure adequate time for the stain to develop on the tissue. After, the slide was washed in an additional coplin jar with Miili-Q water for 3 min to rinse off the excess Cresyl Violet and observe the stain before proceeding with the next rinses. After determining how dark the stain appeared on the tissue, the slide was incubated in 70% ethanol for 1 min and then 95% ethanol for 1 min as well to slightly rinse off excess stain but not entirely remove the stain. The slide was then incubated in xylene for 5 min before being cover slipped with CytoSeal XYL overnight. The slide was imaged the next day on a digital pathology scanner.
j. Choline Acetyltransferase and Cephalotoxin-2. The method was initially developed by Choi et al. (4) with modifications according to Criswell and Gillis (5), and followed here accordingly. Hybridization chain reaction (HCR) reagents were obtained from Molecular Instruments Inc. (LA, USA), where probes for the venom protein cephalotoxin-2 for *D. pealeii* were further specifically designed. The probes for choline acetyltransferase were previously designed for *Octopus bocki* (6), and here tested on *D. pealeii* PSG sections. Sequences upon which the hybridized probes were designed can be found in the supplemental Data S1, while the comparative alignment of the *O. bocki* homolog ChAT sequences from *D. pealeii*, *O. bimaculoides and A. bandense* can be found in the supplementary methods and Fig. S5 while the ChAT cephalopod homolog sequences are found in Dataset S1. The HCR was performed as previously described for the cephalotoxin-1 hybridization in the multiplex experiment (see point e).
k. Live calcium signaling. PSG was dissected from *E. berryi* specimen immediately after euthanasia, and placed in a petri dish with cephalopod-specific cell culture medium, herein called octomedia. Octomedia was prepared as follows (in mM): NaCl: 214.04, MgSO4x7H2O: 26.17, KCl: 4.61, NaHCO3: 2.29, MgCl2x6H2O: 28.04, D-glucose: 38.08 CaCl2x2H2O: 10.12, and 500 ml L-15, Leibowitz, with L-glutamine (Cat.-No. MFCD00217482, Sigma-Aldrich). Octomedia was adjusted to pH 7.6-7.7 with KOH, and filtered using the Corning system (Cat.-No. 431098), stored at 4°C and used within one week. The calcium indicator CAL-520 (Cat.-No. ab171868, Abcam, UK) was prepared at 5 mM in DMSO, and stored at -20°C. CAL-520 improves signal-to-noise ratios compared to standard calcium imaging, allowing live fluorescence recordings of calcium migration within the PSG (7). PSGs were incubated in CAL-520 (5 µM in Octomedia) for 15 min. Imaging was performed with an upright epifluorescent microscope equipped with a 490/525 nm excitation/emission filter set and operated at 2fps. Calcium imaging videos were processed as follows: image registration was performed within suite2p using the non-rigid registration approach (8), spontaneously active regions were manually defined, normalised using a min/max approach and plotted using a custom python script. The maximum intensity projection image shown in Fig. 3 was prepared for publication using ImageJ (9).

***Tissue fixation***

The posterior salivary glands (PSGs) and the digestive gland that aligns anatomical in close proximity to them are both filled with digestive enzymes that can rapidly degrade the tissue samples during extraction (10). Therefore, the fixation of these tissue types have proven to be challenging (11) raising the need of specific protocol adaptations and complex fixation processes. Beside flash freezing and OCT media embedding, the usage of tissue fixative agents is among the most widespread and successful preserving strategies in histology. Among these agents, neutral buffered paraformaldehyde is the most commonly used substance (12). Therefore and following previous studies in cephalopod (*e.g.,* 13), tissue samples were initially fixed in 4% paraformaldehyde (PFA) overnight at 4°C, but at least for 16 h. Afterwards they are subjected to a series of progressive ethanol dehydration steps (20%, 50%, 70%) and tissues are kept in 70% ethanol until needed. This protocol has been challenged throughout our study by the diversity of tissue types and also due to cross-species variability. Thus, following scientific discussion with Dr. Pedro Costa, author of (14) on his experience with the fixation of cuttlefish tissues, Davidson’s/Hartmann’s fixative was hypothesis to have highest potential for tissue fixation success across cephalopods. Davidson’s fixative contains ethanol and acetic acid which is used to penetrate the tissue faster and deeper compared to 4% PFA. Bouin’s fixative was also tested as it contains saturated picric acid, formalin and glacial acetic acid which should also penetrate the tissue rapidly.

The first combinatory fixative trial was conducted due to comparative reasons with available literature background on arm and digestive gland tissue sections of *Octopus bimaculoides,* with additional attempts in *Euprymna berryi* PSG and digestive gland (DG) tissue. Each of the tissue sections was fixed with a different protocol at a minimum of 1:5 tissue to fixative volume ratio: 1) Paraformaldehyde: 4% PFA (1:8 dilution in DEPC-PBS from Cat.No. 15714-S, Electron Microscope Science, Hatfield, USA) overnight incubation, or at least 16 h, at 4°C. 2) Davidson fixative and PFA: Four hours of fixation in Davidson’s/Hartmann’s (Cat.No. 64133-10, Electron Microscope Science) followed by overnight fixation in 4% PFA. 3) Davidson and Bouin fixative: Tissue was fixed in Davidson’s for 4 hours followed by Bouins fixative (Cat. No. 26386-01, Electron Microscope Science) overnight. 4) Bouin and PFA: Tissue was fixed first in 4% PFA followed by overnight fixation in Bouin’s.

The results indicate overall best tissue integrity, stability, and structural diversity preservation success for the Davidson fixative with some exceptions. Davidson/PFA fixative seemed to harden the tissue a lot faster compared to 4% PFA only, however, the *O. bimaculoides* DG tissue was exceptionally soft and sectioning was not possible and thus further verification will be needed. Bouin’s fixative did not appear to accelerate tissue hardening relative to paraformaldehyde, and it resulted in a prominent yellow coloration of the tissues, likely related to the picric acid component. Additional validation is necessary before definitively excluding Bouin’s fixative as a suitable option for this tissue type. Concluding this assessment highlights the need of an adaptive fixative selection in relation to the tissue type and species assessed but also the subsequent analysis to be performed when working in cephalopod histology.

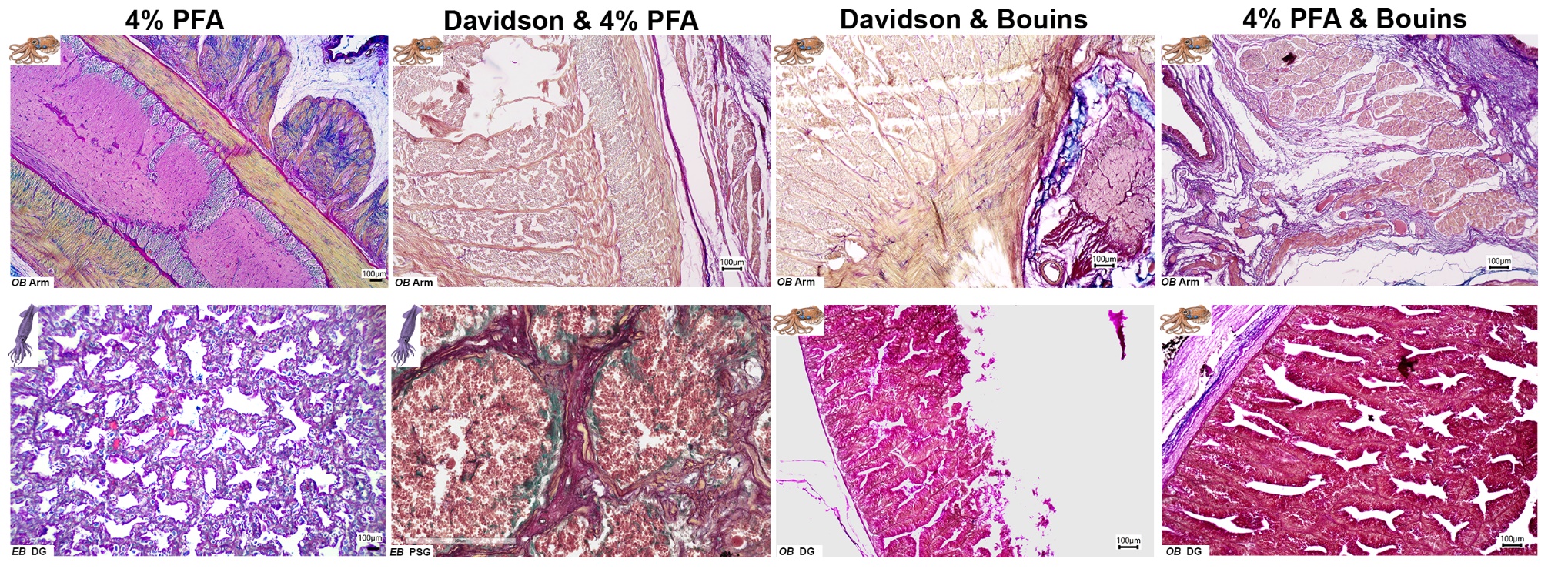

**Figure S4. Tissue fixation requirements vary across tissue types and species.** Comparison between cephalopod tissue samples from Octopus bimaculoides (OB) and Euprymna berryi (EB) fixed with different fixative agents, highlighted on top of the image. Digestive gland (DG) and Posterior salivary gland (PSG), highlight the challenge of digestive enzyme for the tissue preservation.

***Retrieval of cys-loop LGC sequences from the PSG transcriptome***

PSG transcriptomes from *Doryteuthis pealeii* (unpublished dataset and public SRX14223474) *Euprymna berryi* (unpublished dataset), *Ascarosepion bandense* (unpublished data) and *O. bimaculoides* (unpublished dataset and public SRX1045409) were screened herein to identify potential ligand-gated ion channels (LGICs) involved in neuronal control of this glands in cephalopods. Therefore, transcriptomes were analyzed by trimming of the paired end RNA-seq reads to remove adapters and low-quality base calls using Trimmomatic v0.39 or cutadapt v4.2 at default parameters. The success of trimming was assessed with FastQC v0.11.9 (15-17). Sequences for each sample were assembled *de novo* using Trinity v2.15.1 (18).

To identify putative cys-loop LGCs in the PSG of these species, transcriptomes were first analysed using Transdecoder v5.7.1 to identify open reading frames (ORFs) and predict the most probable protein product of each qualifying ORF (19). HMMER v3.3.1 (20) was implemented to scan through all transcriptomes and identify proteins containing the neurotransmitter-gated ion-channel ligand binding domain described in Pfam profile PF02931 (21). Sequences that passed HMMER scanning were funnelled into InterProScan v5.72 (22) to confirm a domain match to PF02931. All sequences that survived the above filtering steps were then analyzed with TMHMM2 to identify transmembrane helices within the proteins (23,24). Candidate cys-loop LGICs that had four transmembrane domains identified via TMHMM2 were passed onto phylogenetic analysis. The results of this search are given in the supplemental Data S1, and it must be noted that LGIC sequences with the corresponding domain match could not be confirmed for *E. berryi*.

***Choline acetyltransferase cross-cephalopod-species sequence alignments – homology***

Acetylcholine is a dominant neurotransmitter across multiple metazoan lineages, as recovered in the neuroreceptor phylogenetic analysis presented in the main manuscript (Fig. 4). Based on this finding, we targeted choline acetyltransferase (ChAT), the enzyme responsible for acetylcholine synthesis, in the posterior salivary gland (PSG) of cephalopods in order to characterize its abundance and spatial distribution within the gland.

A previously designed ChAT RNA probe developed for hybridization chain reaction (HCR) in *Octopus bocki* (6) was used as a query in BLAST searches against transcriptomes of *D. pealeii*, *O. bimaculoides*, and *A. bandense* to assess sequence conservation and evaluate the feasibility of cross-species probe hybridization. This analysis identified a total of 95 transcript matches across the three species, of which 92 exhibited moderate sequence identity to the *O. bocki* ChAT probe (25–60%). Three transcripts with high sequence similarity (>95% identity) were identified in *D. pealeii* and *O. bimaculoides* (supplemental Data S1). Based on these results, ChAT HCR experiments were conducted in these two species. For species-specific alignments of the ChAT homologs retrieved from the transcriptomes see Figure S5.

The design of species-specific ChAT probes for *E. berryi* and *A. bandense* is underway and will allow expansion of this comparative framework, enabling a broader assessment of acetylcholine metabolism and cholinergic signaling pathways in the cephalopod PSG.

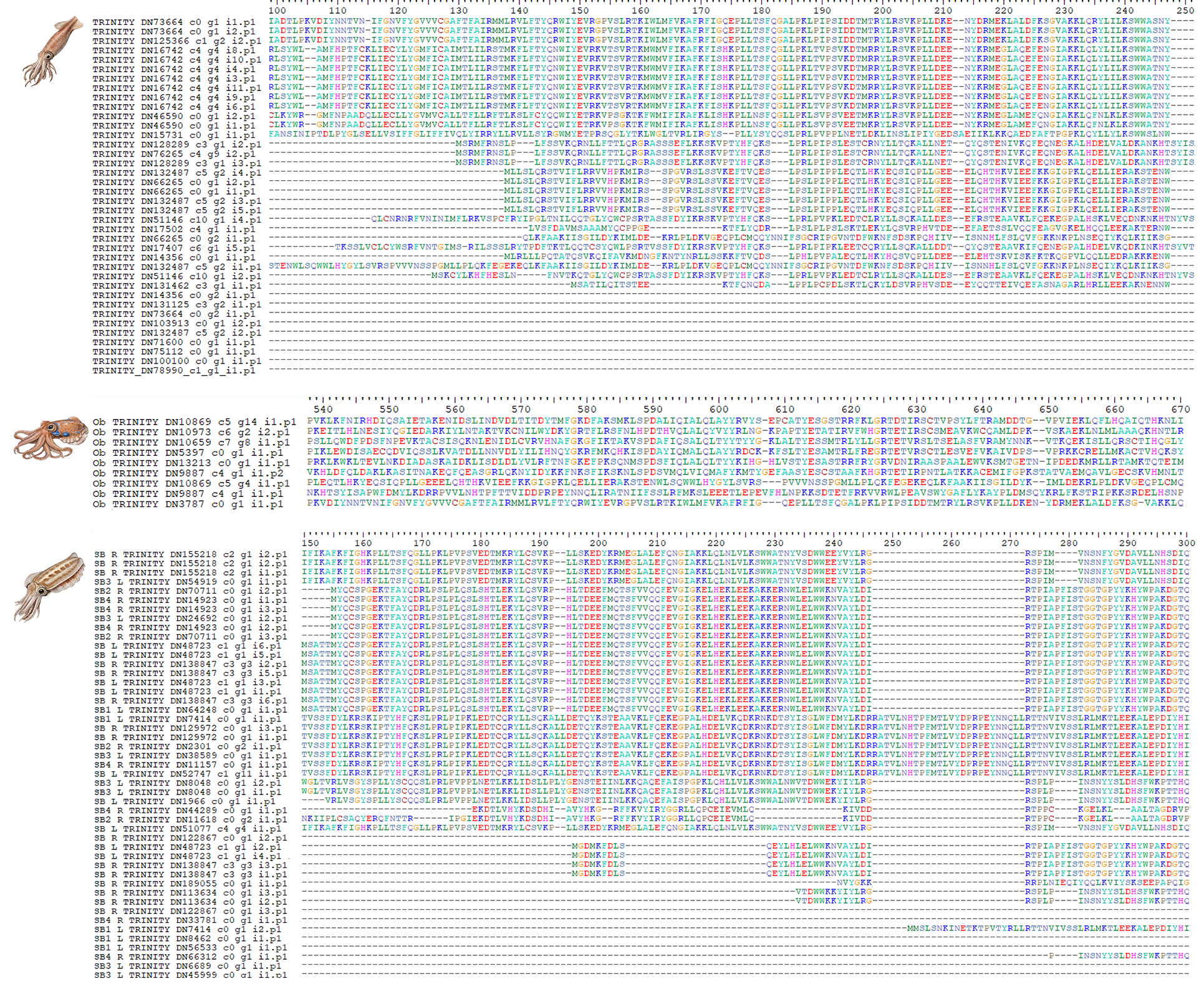

**Figure S5.** Choline acetyltransferase (ChAT) transcript sequence alignments from *Doryteuthis pealeii*, *Octopus bimaculoides* and *Ascarosepion bandense*. These sequences have been retrieved from the posterior salivary gland transcriptomes by a BLAST search with the *Octopus bocki* ChAT sequence as query. The sequence were translated to amino acid sequences for similarity comparison.

Movie S1.

Maximum intensity projection recorded in *ex vivo* posterior salivary glands of adult *Euprymna berryi* during calcium imaging.

Data S1. (separate file)

The dataset includes the original sequences of *Octopus bocki* and *Doryteuthis pealeii* upon which the probes for the hybridization chain reaction, targeting choline acetyltransferase and cephalotoxin-2, were designed. Further, we include the information of the amplifier and fluorophores employed.
